# Integrative Modeling and Analysis of Fungal Central Carbon Metabolism

**DOI:** 10.64898/2026.05.08.723079

**Authors:** Janaka N. Edirisinghe, Claudia Lerma-Ortiz, Filipe Liu, Jose Faria, Robert W. Cottingham, Adam P. Arkin, Qun Liu, Christopher S. Henry

## Abstract

Over a thousand fungal genomes have been sequenced, yet manually curated genome-scale metabolic models (GEMs) are available for only a limited number of species. Moreover, these models have often been developed independently, leading to inconsistencies in namespaces, compartment definitions, and pathway representations that hinder comparative analysis, the systematic reuse of prior curation efforts, and the integration of consolidated metabolic knowledge. Here, we present the Consolidated Fungal Core Metabolism Model (CFCMM), constructed by integrating thirteen published fungal models spanning Ascomycota, Mucoromycota, and both Crabtree-positive and Crabtree-negative yeasts. We harmonized metabolites and reactions into a non-redundant shared ModelSEED ontological space, standardized compartmentalization, and refined gene–protein–reaction (GPR) rules. Using pathway-level visualization and systematic gap detection, we further improved the integrated network through literature-guided curation to correct stoichiometry, stereospecificity, and pathway architecture. Orthologous protein family reconstruction and functional annotation workflows were used to validate and inform GPR associations, with particular emphasis on ambiguous enzyme superfamilies and membrane-associated components.

Using the resulting CFCMM, we built high-quality central carbon core models for each fungus and performed flux balance analysis to quantify ATP-yield variation under aerobic and anaerobic conditions, explicitly evaluating scenarios driven by differences in electron transport chain (ETC) composition. Simulations reproduced the expected fermentative yield of approximately 2 mmol ATP per mmol glucose under anaerobic conditions and separated the thirteen fungi into two bioenergetic groups under aerobic respiration based on Complex I status, with predicted yields of approximately 30 versus 22 mmol ATP per mmol glucose. Forcing flux through the alternative oxidase bypass further reduced ATP yields to approximately 12 and 4 mmol ATP per mmol glucose in Complex I-containing and Complex I-lacking fungi, respectively. Collectively, this work provides a manually curated, ModelSEED-consistent, and extensible fungal core metabolic template, deployed in DOE KBase as a resource for automated reconstruction of central carbon core models from any sequenced fungal genome. In addition, the CFCMM provides modular components for developing GEMs with more accurate energy predictions and enables robust comparative analyses of fungal bioenergetics and core metabolic diversity.

## Introduction

Fungi are major actors in biogeochemical cycling, agriculture, industrial biotechnology, and human disease, and their metabolic capabilities are as diverse as the habitats they occupy. A deeper understanding of fungal biology is therefore essential for advancing biotechnology, addressing challenges in health and agriculture, and promoting environmental sustainability (Table S1). Consistent with this broad ecological and applied importance, fungi have evolved to thrive in diverse environments, including soil and marine ecosystems, and under a wide range of physicochemical conditions such as varying pH, temperature, and oxygen availability.

Despite their ecological and industrial importance, our understanding of fungal biology remains limited given the immense diversity of the fungal kingdom. Over the past decade, advances in genomics and systems biology have enabled large-scale sequencing efforts and the application of *in silico* metabolic modeling to investigate fungal physiology, biochemistry, and behavior. To date, more than a thousand fungal genomes have been sequenced and annotated (Kim et al., 2023; *1000 Fungal Genomes Project*, n.d.), and several genome-scale metabolic models have been developed (Brandl & Andersen, 2015).

Furthermore, metabolic models play a critical role in studying fungi across diverse contexts, including pathogenic systems and fungal interactions with plant hosts and microbial communities. While automated reconstruction of metabolic models for prokaryotes is supported by several well-established tools, the development of manually-curated, biochemically-consistent models for individual fungi remains challenging. This is largely due to limitations in genome annotation and the incomplete representation of fungal-specific biochemistry in existing databases. Although a limited number of published fungal models provide curated biochemical knowledge, there is a clear need to reconcile these datasets within a unified ontological framework. Such integration would enable the development of a standardized template for fungal metabolism, thereby facilitating the generation of draft metabolic models for newly sequenced genomes.

In this context, our study focuses on fungal central carbon metabolism, as it is fundamental for accurately capturing energy-biosynthesis strategies and predicting energy yields. This approach is analogous to previous work in prokaryotic systems (Edirisinghe et al., 2018, 2016). Prior studies have shown that energy yields derived from core central metabolism are essential for reliable model predictions and for accurately representing their effects on growth (Faria et al., 2023; Henry et al., 2010).

Establishing a consolidated framework for fungal central metabolism enables several key advances. First, it facilitates the automated reconstruction of core fungal metabolic models and supports comparative, systems-level analyses of metabolic functions across diverse fungal species (analysis in progress). Second, it enables the systematic characterization of metabolic diversity and unique biochemical capabilities while providing a robust foundation in the form of a Consolidated Fungal Core Metabolism Model (CFCMM), which can be extended toward genome-scale metabolic reconstructions. Third, it enhances the ability to model fungal interactions within microbial communities, improving our understanding of their ecological roles and functional dynamics.

With this objective in mind, here we consolidate thirteen published fungal genome-scale metabolic models (Table 1), spanning Ascomycota and Mucoromycota, into the ModelSEED ontology (Seaver et al., 2021) and systematically evaluate the biochemical pathways and reactions associated with central metabolism. Pathways extracted from the published models were reconciled into a non-redundant biochemical representation and further refined through gap detection, literature-anchored reaction chemistry, phylogenetically informed ortholog families, and DRAM-based functional annotation, with particular care given to enzyme superfamilies and membrane-associated components for which sequence homology alone can be misleading. The resulting integration produced a unified Consolidated Fungal Core Metabolism Model (CFCMM). Using this framework, we generated species-specific central carbon core models for thirteen fungi and used flux balance analysis (FBA) to quantify energy-yield variation across electron transport chain (ETC) architectures under aerobic and anaerobic conditions. We further resolved annotation gaps and missing or incorrect biochemistry, thereby improving biochemical consistency and model predictive capability. The CFCMM and its thirteen species-specific core models are released in KBase as a template for automated reconstruction of core metabolic models from any sequenced fungal genome, providing a resource for comparative analysis of fungal bioenergetics and core metabolic diversity.

**Table 1.** Published fungal metabolic models used for construction of the CFCMM. The thirteen published fungal models used to derive initial biochemical content, gene–protein–reaction associations, compartmentalization information, electron transport chain composition, and pathway structures for construction of the CFCMM, as depicted in Fig. 1.

| Model Id | Organism | Phylum | Features | Reactions | Crabtree Positive/Negative | Reference |
| --- | --- | --- | --- | --- | --- | --- |
| iJL1454 | *Aspergillus terreus* NIH2624 | Ascomycota | 1431 | 1276 | mold; not typically classified | [doi: 10.1039/c3mb70090a](https://doi.org/10.1039/c3mb70090a) |
| iNX804 | *Candida glabrata* ASM254 | Ascomycota | 804 | 1179 | Positive | [doi: 10.1039/c2mb25311a](https://doi.org/10.1039/c2mb25311a) |
| iCT646 | *Candida tropicalis* MYA-3404 | Ascomycota | 645 | 1265 | Negative | [doi: 10.1002/bit.25955](https://doi.org/10.1002/bit.25955) |
| iRL766 | *Eremothecium gossypii* ATCC 10895 | Ascomycota | 752 | 1430 | Not consistently labeled in the literature | [doi: 10.1002/bit.25167](https://doi.org/10.1002/bit.25167) |
| iOD907 | *Kluyveromyces lactis* NRRL | Ascomycota | 911 | 1866 | Negative | [doi: 10.1002/biot.201300242](https://doi.org/10.1002/biot.201300242) |
| iLC915 | *Komagataella phaffii* GS115 | Ascomycota | 914 | 1389 | Negative | [doi: 10.1186/1752-0509-6-24](https://doi.org/10.1186/1752-0509-6-24) |
| iJDZ836 | *Neurospora crassa* OR74A | Ascomycota | 828 | 1360 | mold; not typically classified | [doi: 10.1371/journal.pcbi.1003126](https://doi.org/10.1371/journal.pcbi.1003126) |
| iAL1006 | *Penicillium chrysogenum* Wisconsin | Ascomycota | 995 | 1465 | mold; not typically classified | [doi: 10.1371/journal.pcbi.1002980](https://doi.org/10.1371/journal.pcbi.1002980) |
| iSS884 | *Scheffersomyces stipitis* CBS | Ascomycota | 883 | 1320 | Negative | [doi: 10.1186/1752-0509-6-24](https://doi.org/10.1186/1752-0509-6-24) |
| iNL895 | *Yarrowia lipolytica* CLIB122 | Ascomycota | 898 | 1799 | Negative | [doi: 10.1186/1752-0509-6-35](https://doi.org/10.1186/1752-0509-6-35) |
| iWV1213 | *Mucor circinelloides* CBS277 | Mucoromycota | 1212 | 1249 | Positive | [doi: 10.1016/j.gene.2016.02.028](https://doi.org/10.1016/j.gene.2016.02.028) |
| iWV1314 | *Aspergillus oryzae* RIB40 | Ascomycota | 1346 | 1248 | mold; not typically classified | [doi: 10.1186/1471-2164-9-245](https://doi.org/10.1186/1471-2164-9-245) |
| iMM904 | *Saccharomyces cerevisiae* 5288c | Ascomycota | 905 | 1412 | Positive | [doi: 10.1089/ind.2013.0013](https://doi.org/10.1089/ind.2013.0013) |

## Methods

Supplementary Tables can be accessed via Figshare at: https://doi.org/10.6084/m9.figshare.32114788 Fungal core metabolic model reconstructions - KBase Narrative https://narrative.kbase.us/narrative/251530 (static Narrative - https://kbase.us/n/251530/77/)

Our overarching goal is to assemble a complete and biochemically consistent set of core pathways that together form a consolidated metabolic model, which can serve as a template for reconstructing central carbon core metabolic models for any fungal genome. Fig 1 outlines the workflow used to construct the Consolidated Fungal Core Metabolism Model (CFCMM). The methods used in each step of this workflow are described below.

**Fig 1.**
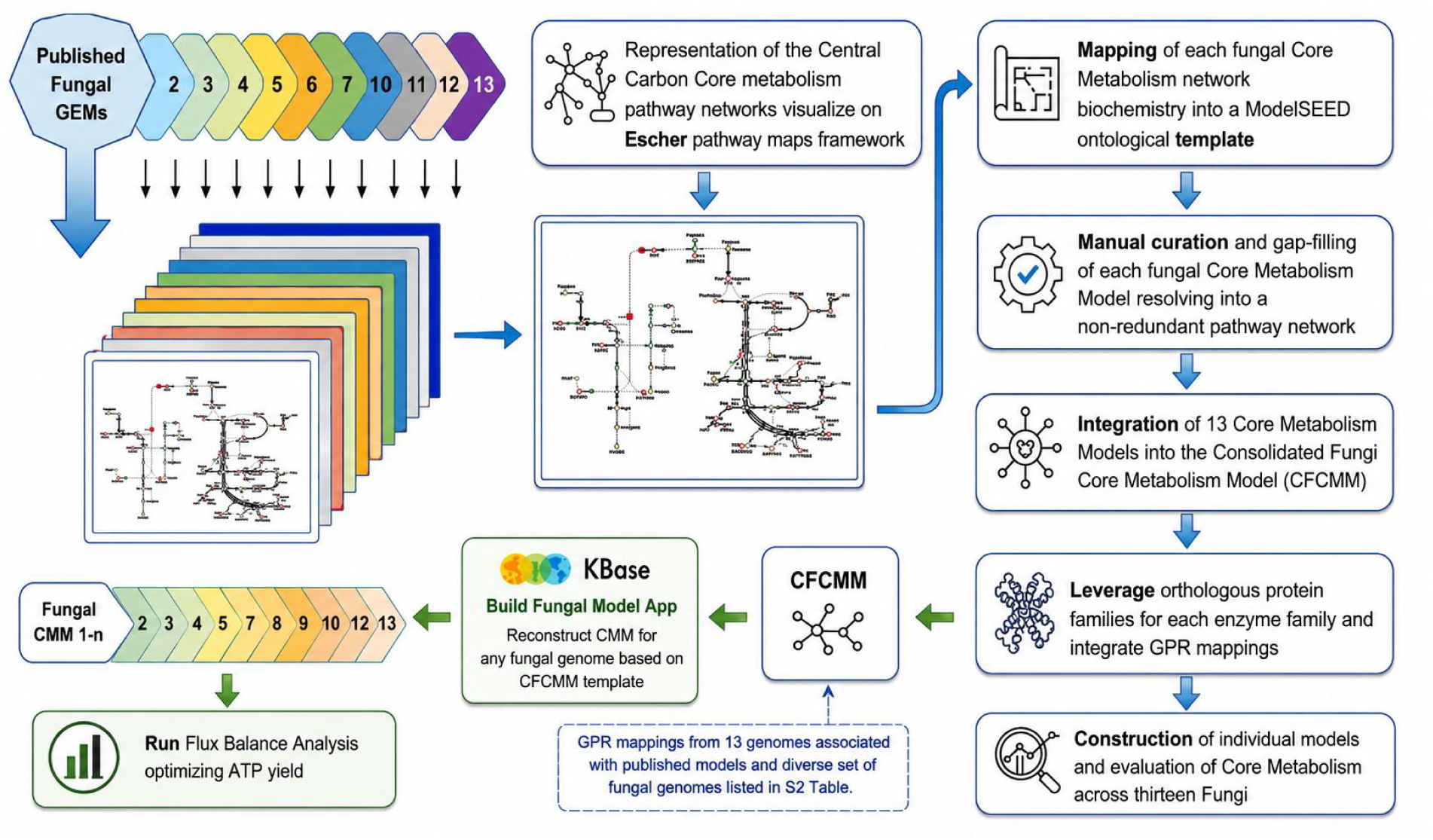
Construction pipeline for the Consolidated Fungi Core Metabolism Model (CFCMM). Biochemical representations from the 13 published models listed in Table 1 were used to construct the CFCMM. Central carbon core metabolism networks from these models were evaluated and visualized using the Escher pathway-mapping framework, and their biochemistry was reconciled within the ModelSEED ontology, then extensively manually curated to improve consistency across models, reduce redundancy, and facilitate gap detection. In parallel, orthologous protein families were computed from the fungal genomes underlying the published models, together with the additional fungal genomes listed in S2 Table, thereby expanding enzyme-family mapping and supporting the integration of gene–protein–reaction (GPR) associations into the CFCMM. The curated CFCMM was then used as a template for reconstructing central carbon core metabolism models for any fungal genome through the Build Fungal Model app in KBase, followed by flux balance analysis to evaluate ATP yield, as summarized in Table 2

**Table 2.**
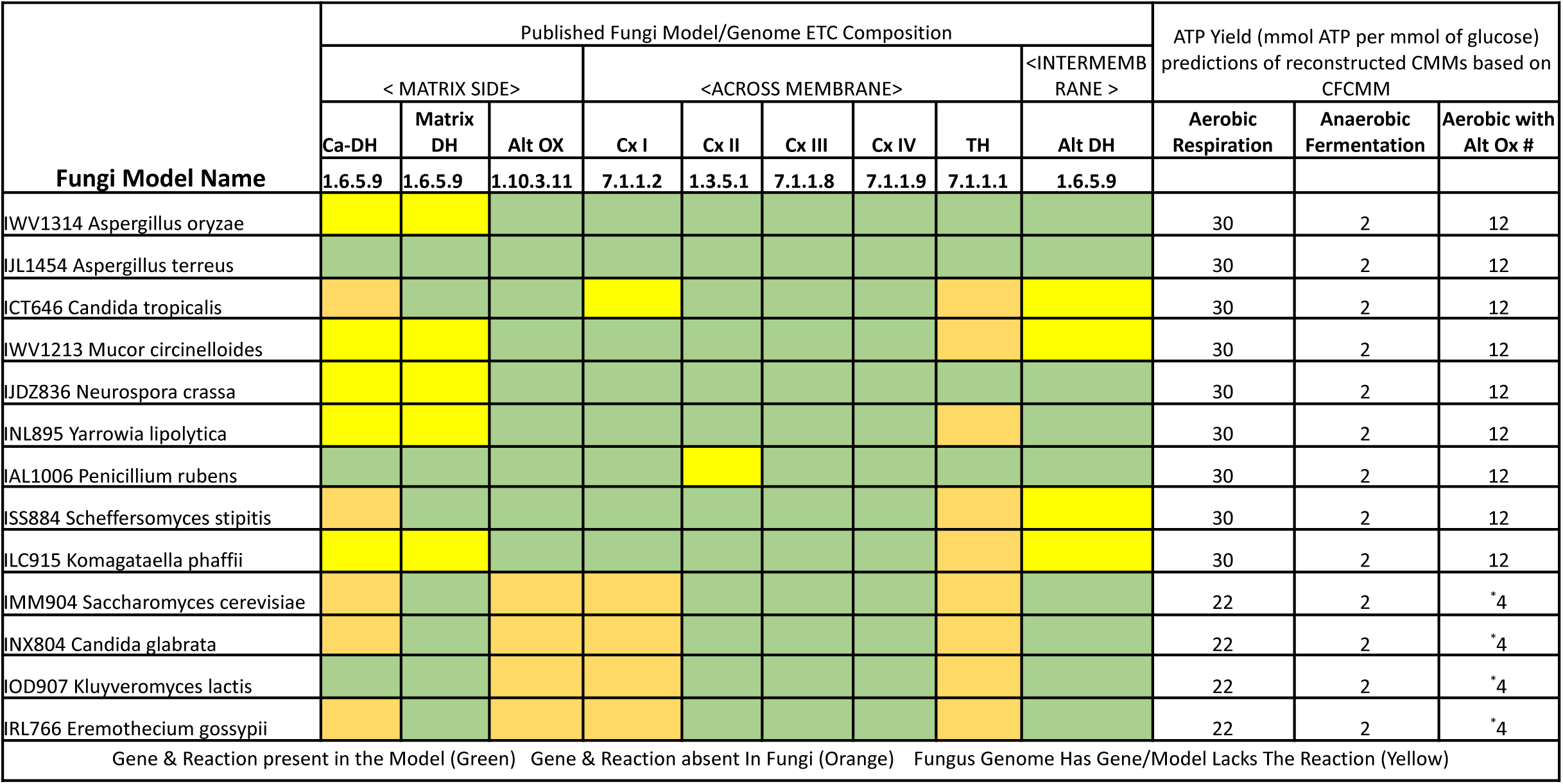
Relationship between ATP-yield predictions in CMMs and the enzymatic composition of the fungal mitochondrial inner membrane. ATP-yield predictions were generated by flux balance analysis (FBA) under aerobic and anaerobic conditions, using glucose as the sole carbon source, for each reconstructed fungal core model based on the CFCMM (white columns). Colored columns indicate the Enzyme Commission (EC) numbers of electron transport chain (ETC) components and the presence or absence of the corresponding enzymes in the published models of each fungus (see row above for color definitions). Where the original published models lacked a reaction but the corresponding genes were present in the genome, those reactions and GPR mappings were added to the CFCMM. ATP yields calculated after blocking flux through Complex III and Complex IV (#*) represent the hypothetical yields expected in the presence of an active alternative oxidase in the mitochondrial electron transport chain. All reconstructed models and FBA simulations showing key variations on ATP-yields are available in the KBase Narrative at https://narrative.kbase.us/narrative/251530 (https://kbase.us/n/251530/77/). Abbreviations: Ca-DH, calcium-dependent NADH dehydrogenase; Matrix DH, matrix NADH dehydrogenase; Alt OX, alternative oxidase; Cx, complex; TH, transhydrogenase; Alt DH, alternative NADH dehydrogenase.

### Conversion of GEMs

We surveyed the literature and selected 13 diverse published fungal genome-scale metabolic models, from which we extracted curated biochemical pathways, compartmentalization data, and gene–protein–reaction (GPR) associations (Table 1). Together, these models encompass both Crabtree-positive and Crabtree-negative fungi (De Deken, 1966) (Table 1, S1 Table). We converted these GEMs into networks operating within the ModelSEED biochemistry framework (Seaver et al., 2021)(https://modelseed.org/) by reconciling diverse biochemical nomenclatures into a non-redundant representation and linking the resulting reactions to their corresponding GPR relationships.

### Construction of core metabolism (CoM) networks

To construct the CoM networks, we selected key central carbon metabolic pathways from 13 genome-scale metabolic models (GEMs) that capture variations in energy-biosynthesis strategies and sugar metabolism. These pathways include glycolysis/gluconeogenesis, the pentose phosphate pathway (PPP), the tricarboxylic acid (TCA) cycle, the glyoxylate cycle (GlxC), the γ-aminobutyric acid (GABA) shunt, diverse fermentation pathways, and variations in the fungal aerobic respiratory electron transport chain (ETC).

For gap detection and evaluation of the reconciled non-redundant biochemistry, each CoM network was graphically represented using the Escher visualization environment (King et al., 2015), with the yeast model iMM904 (Table 1) used as the starting point because it is the most comprehensively resolved within the ModelSEED ontology and is also among the simplest of the models examined. We then performed detailed manual curation to identify and resolve pathway gaps. Complementing the biochemical curation, we computed orthologous protein families associated with each enzyme class across the fungal genomes from which the published models were derived using the OrthoMCL algorithm (L. Li et al., 2003) deployed in the KBase platform (Arkin et al., 2018). In addition, we expanded these protein families by incorporating sequences from an additional 15 fungal genomes (S2 Table), enabling the capture and reconciliation of biochemical functions that extend phylogenetically and metabolically beyond the genomes represented in the published models. Furthermore, all fungal genomes were annotated using the DRAM functional annotation pipeline (Shaffer et al., 2020, 2023), which assigns Enzyme Commission (EC) numbers and KEGG Orthology (KO) identifiers to predicted proteins. These annotations were evaluated to verify the accuracy of associated enzymatic reactions and to assess consistency across protein families mapped to the same biochemical functions. The DRAM annotations for all fungal genomes are available in the associated KBase Narrative https://narrative.kbase.us/narrative/251530 (static Narrative - https://kbase.us/n/251530/77/).

Finally, using the curated biochemical reactions and GPR associations supported by orthologous protein families, we constructed a highly curated consolidated metabolic network template representing fungal central carbon metabolism. This template was subsequently used to build central carbon core metabolic models based on individual fungal genomes.

### Using Build Fungal Model App in KBase

CoM models were constructed using the DOE KBase fungal model reconstruction app “Build Fungal Model” (https://narrative.kbase.us/legacy/catalog/modules/kb_fungalmodeling) (Arkin et al., 2018; Edirisinghe et al., n.d.) starting from a consolidated metabolic network that integrates all curated biochemical reactions and associated GPR rules. The app accepts any structurally annotated fungal genome as input and infers orthologs using a bidirectional best BLAST hit (BBH) approach against the user-provided genome to assign GPRs. Based on these mappings, the app propagates the relevant biochemical reactions from the consolidated metabolic network template to generate a draft core model for the genome of interest. The process for building a core metabolic model is demonstrated in the KBase Narrative at https://narrative.kbase.us/narrative/251530 (https://kbase.us/n/251530/77/). Furthermore, all thirteen curated central carbon core metabolic models generated in this study are available in the same Narrative.

### Validation of the CFCMM by functional analysis

For the central carbon core metabolism models, ATP hydrolysis (H₂O + ATP → ADP + phosphate + H⁺ - in the cytosolic compartment) was used as the objective function, reflecting the primary goal of accurately predicting energy yields through flux balance analysis (FBA). *In silico* simulations were performed using a glucose minimal medium in which glucose served as the sole carbon source under both aerobic and anaerobic conditions. Model simulations were conducted using FBA implemented through both the modelseedpy package (https://github.com/ModelSEED/ModelSEEDpy) and the FBA app available within the DOE KBase platform (https://narrative.kbase.us/legacy/appcatalog/app/fba_tools/run_flux_balance_analysis/release). To evaluate energy yields while capturing variations in fungal electron transport chains, simulations were performed with the glucose uptake rate constrained to 1 mmol gDW⁻¹ h⁻¹ under both aerobic and anaerobic conditions. ATP production rates were then calculated to assess model-predicted energy yields (Table 2).

## Results and discussion

### ModelSEED representations of fungal core metabolism networks and their mapping

As described in the Methods section, we used a custom-tailored Escher map environment to reconcile fungal GEMs with the ModelSEED ontological framework. S3 Table summarizes the analysis of these mapped networks, highlighting pathway gaps and the corresponding corrections proposed for each model. Several recurring issues were identified across the reactions, including inconsistencies in stereoisomer specificity, reaction stoichiometry, and substrate specificity. Addressing these inconsistencies and establishing consistent solution patterns substantially simplified the gap-filling process (S4 Table) (see the section on CFCMM construction following the manual curation section for additional details).

After implementing these initial reaction-level corrections, it became evident that a more comprehensive curation of pathway architecture and functional integration across central metabolic pathways was required. This need arises because existing fungal GEMs often emphasize curation of specific metabolic subsystems depending on the particular biological or biotechnological application for which the model was developed, rather than focusing on a consistently curated accurate representation of core metabolism.

### Evidence based manual curation and gap filling

In this section, we describe the evidence-driven decision process used to identify, correct, and fill gaps in core metabolism. We also highlight key pathways and discuss how their biological roles shape the organization and behavior of the integrated metabolic network.

### Mitochondrial NADH dehydrogenases, their location and calcium-controlled activity

#### Role

The duplication of alternative NADHDHs (both internal and external) coincides with the loss of Complex I in the *Saccharomyces* and *Kluyveromyces* lineages—potentially enhancing NADH reoxidation and sustaining NAD⁺ regeneration under fermentative conditions (Marcet-Houben et al., 2009). Gene duplication patterns further support functional diversification of these enzymes. In addition, a distinct subset of NADHDHs contains an EF-hand calcium-binding motif; Notably, EF-hand NADHDHs are more frequently observed in fungi that retain Complex I, whereas fungi lacking Complex I often do not encode the EF-hand form. This pattern suggests that calcium-dependent control of NADH oxidation may be preferentially maintained in Complex I–positive respiratory architectures, consistent with the broader role of mitochondrial calcium signaling in regulating respiratory function (Del Arco et al., 2016; McDonald & Gospodaryov, 2019).

#### Curation

Because mitochondrial NADH dehydrogenases (NADHDHs) often have high degree of sequence similarity and typically carry mitochondrial targeting peptides, sequence features alone are insufficient to reliably infer membrane orientation; definitive assignment generally requires experimental validation. Membrane localization and orientation have been characterized for *Saccharomyces cerevisiae* (Luttik et al., 1998; Small & McAlister-Henn, 1998; Marres et al., 1991), *Yarrowia lipolytica* (Kerscher et al., 1999), and *Neurospora crassa* (Duarte et al., 2003; Weiss et al., 1970). In *N. crassa*, the EF-hand calcium-binding motif containing enzyme was shown to be internally oriented (matrix-facing) based on detergent accessibility and alkaline extraction assays (Melo et al., 2001). Consistent with these experimental observations, our phylogenetic reconstruction of fungal NADHDHs (S1 Fig) separates matrix-facing enzymes (non–Complex I), from alternative NADHDHs in both *S. cerevisiae* and *N. crassa*, and places EF-hand–containing NADHDHs into a single clade that includes the *N. crassa* EF-hand enzyme. Similar relationships between phylogenetic grouping and membrane orientation have been noted previously (Kerscher et al., 1999; Marcet-Houben et al., 2009). Most fungi examined here encode more than one mitochondrial NADH dehydrogenase, although EF-hand–containing forms are not universally present (S5 Table). This pattern is biologically plausible, though it will require targeted experimental confirmation before being used to assign orientation in newly reconstructed models. For the CFCMM, we therefore adopted a conservative strategy: we assume the presence of at least one matrix-facing NADHDH across taxa, regardless of Complex I status, and include this activity in the consolidated template.

#### NADH shuttles

The inner mitochondrial membrane is impermeable to NADH, and cells regulate NADH/NAD homeostasis primarily through three shuttles to transport NADH across this membrane: the ethanol-acetaldehyde shuttle (EAS), the aspartate-malate shuttle (AMS), and the glycerol 3-phosphate shuttle (G3PS) (Fig 2 and S6 Table). Some of them are linked indirectly to ETC flow.

**Fig 2.**
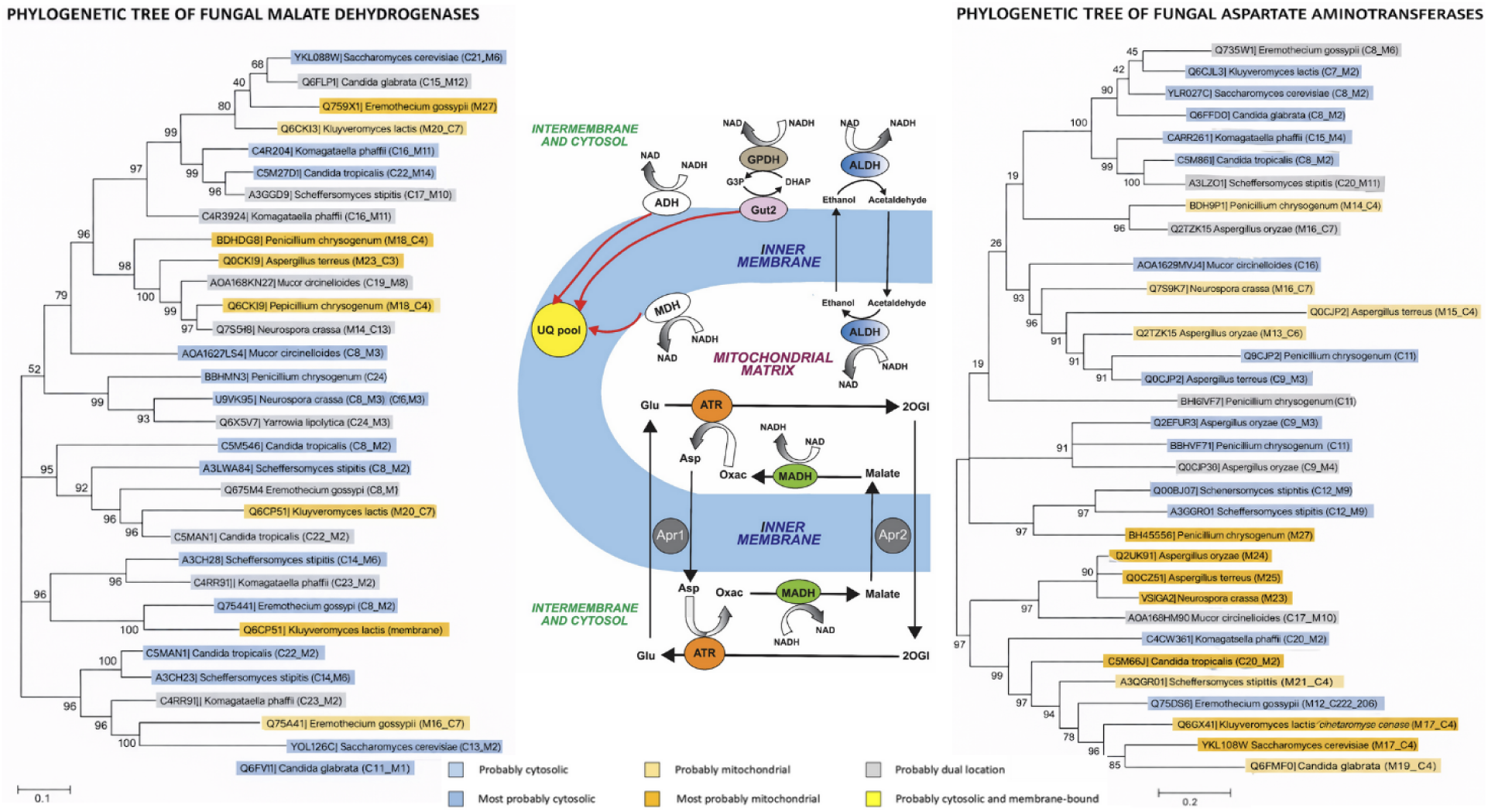
NADH shuttles and protein-based phylogenetic trees of aspartate aminotransferases and malate dehydrogenases.

### Ethanol–acetaldehyde shuttle

#### Role

The EAS provides an important mechanism for redox balancing between the cytosol and mitochondria, particularly in fungal species lacking matrix NADH dehydrogenases or Complex I—key components typically responsible for transferring electrons from mitochondrial NADH to the ubiquinone pool. In the absence of these, the EAS offers an alternative route to channel electrons into the ETC and maintain energy production. In this shuttle, NADH is oxidized to NAD⁺ as acetaldehyde is reduced to ethanol within the mitochondrial matrix. The produced ethanol diffuses across the membrane into the cytosol, where it is re-oxidized to acetaldehyde, coupled with the reduction of cytosolic NAD⁺ to NADH. This cytosolic NADH can then transfer its electrons to the ubiquinone pool via an alternative NADH dehydrogenase localized on the cytosolic face of the inner mitochondrial membrane(Luttik et al., 1998; Merico et al., 2007) (Fig 2). The EAS is reversible and can also operate in the opposite direction. When cytosolic NADH accumulates—for instance, in the absence of cytosolic NADH dehydrogenase and the glycerol-3-phosphate shuttle (see Function of the Glycerol-3-Phosphate Shuttle section)—acetaldehyde is reduced to ethanol in the cytosol, which then diffuses into the mitochondrion. There, ethanol is oxidized back to acetaldehyde, regenerating mitochondrial NADH that can feed electrons into the ETC via any remaining matrix dehydrogenase (Bakker et al., 2000; Boubekeur et al., 2002). This flexibility makes the EAS an important compensatory mechanism for fungi lacking canonical ETC components, as is observed in *Saccharomyces cerevisiae* and other yeast species.

#### Curation

Enzymes catalyzing the alcohol dehydrogenase reaction are present in both cytosolic and mitochondrial compartments in all the studied models (S6 Table), except for *Neurospora crassa.* However, we identified NCU02476, a gene encoding for this enzyme, and predicted it to be mitochondrial, which would enable it to have the EAS.

The ethanol-acetaldehyde, aspartate-malate and glycerol 3-phosphate shuttles indirectly transport NADH across the mitochondrial inner membrane. They do not operate at the same time, nor are all of them present in all fungi. Abbreviations: MDH, matrix NADH dehydrogenase; ADH, alternative NADH dehydrogenase; UQ, ubiquinone; The reaction directions shown in the figure are representative of the following mechanisms: **for the aspartate-malate shuttle** the net transport shown is NADH[c0] => NADH[m0]; Abbreviations: ATR, aspartate aminotransferase; Glu, L-glutamate; 2OGl, 2-Oxoglutarate; Asp, L-aspartate; Oxac, oxaloacetate; Apr1, L-glutamate/ L-aspartate antiporter; Apr2, L-malate/2-oxoglutarate antiporter; MADH, malate dehydrogenase. **For the glycerol 3-phosphate shuttle**, the direction indicates the electron donation from NADH[c0] into the ubiquinone pool[m0]; Abbreviations: GPDH, cytosolic NAD dependent glycerol 3-phosphate dehydrogenase; Gut2, mitochondrial FAD-dependent glycerol 3-phosphate dehydrogenase; G3P, glycerol 3-phosphate; DHAP, dihydroxyacetone phosphate; This transfer of electrons is equivalent to the NADH being transported into the matrix and being used by the MDH, which would in its turn donate electrons to the ubiquinone pool**. For the ethanol-acetaldehyde shuttle,** the reactions’ direction represents the mechanism for the <u>NADH</u> expulsion out of the matrix into the cytosol, when there is an NADH excess inside the mitochondria. If this shuttle is used for the introduction of NADH into the mitochondria, then the reactions will all have the opposite directions to those shown in this figure. Abbreviations: ALDH, alcohol dehydrogenase.

### Glycerol-3-phosphate shuttle

#### Role

The G3PS shuttle in fungi is a key mechanism for transferring electrons from cytosolic NADH into the mitochondrial ETC, particularly in species lacking Complex I and ADH (Fig 2). In our Consolidated Fungal Core Metabolism Model we have integrated and simulated the function of the G3PS (See corresponding section). During glycolysis, cytosolic NAD⁺ is reduced to NADH, but because NADH cannot cross the mitochondrial membrane, the G3PS enables its reoxidation, which is essential for sustaining glycolytic flux (de Vries & Marres, 1987). In this process, cytosolic glycerol-3-phosphate dehydrogenase (GPD) reduces dihydroxyacetone phosphate (DHAP) to G3P using NADH. Gut2, a mitochondrial G3P dehydrogenase embedded in the inner membrane, reduces FAD to FADH₂ in the process. The resulting FADH₂ donates electrons to the ubiquinone pool, bypassing Complex I (Shi et al., 2018). This mechanism is particularly important in fungi such as *Saccharomyces cerevisiae*, *Candida glabrata* and *Eremothecium gossypii* which have evolved ETC architectures that lack Complex I (Marcet-Houben et al., 2009; Merico et al., 2007). Although bypassing Complex I results in a lower ATP yield per NADH (see energy yield predictions in Table 2), the G3P shuttle remains an essential pathway for enabling oxidative phosphorylation when other routes are absent (Schertl & Braun, 2014).

#### Curation

A third reaction involved in the G3PS is the oxidation of the FADH2 by ubiquinone, hence connecting the shuttle to its ETC. In the 13 models there was a wide variety in the content of the G3PS three constituting reactions (S6 Table), so we designed a uniform set of them to represent the G3PS in all 13 fungi and to facilitate the integration process needed for the consolidated model. We also identified two *Mucor circinelloides* FAD dependent glycerol-3-phosphate dehydrogenases (OAD02414.1; OAD02109.1), adding their corresponding reactions to this model and enabling its G3PS.

### Malate–aspartate shuttle

#### Role

The MAS also facilitates the transfer of electrons from cytosolic NADH into the mitochondrial matrix, enabling the continuation of glycolysis and promoting efficient ATP production through oxidative phosphorylation. MAS serves as an indirect yet highly efficient mechanism to maintain redox balance by regenerating cytosolic NAD⁺. In this shuttle, cytosolic malate dehydrogenase reduces oxaloacetate to malate using NADH. Malate is then transported into the mitochondrial matrix, where it is re-oxidized by malate dehydrogenase, producing NADH that transfers electrons to Complex I (Spinelli & Haigis, 2018) (Fig 2). This shuttle enables fungi that retain Complex I—such as A*spergillus nidulans* and *Neurospora crassa*—to achieve higher ATP yields by transferring electrons to the ubiquinone pool in the ETC. However, organisms that have the ADH in their ETC, or a functional G3PS or EAS are unlikely to utilize the MAS as those are convenient direct cofactor consumers. The AMS relies on the Apr1 antiporter (L-glutamate/L-aspartate), which depends on the electrochemical potential (Borst, 2020).

#### Curation

At first sight, most of the reactions needed to have a functional MAS in the 13 models were already present, except for the *Neurospora crassa* model which was missing the mitochondrial L-aspartate:2-oxoglutarate aminotransferase reaction. Since we identified NCU08411 as this enzyme, we added this reaction to the model. Fungi have multiple malate dehydrogenase isoforms (S6 Table). *Saccharomyces cerevisiae* has two malate dehydrogenases Mdh1p (UniProt ID: P17505, AF_AFP17505F1) and Mdh2p (UniProt ID: P22133, AF_AFP22133F1) which are mainly cytosolic, according to location prediction programs like WoLF PSORT. *S. cerevisiae’s* third malate dehydrogenase, Mdh3p (UniProt ID: P32419, AF_AFP32419F1), is predicted as peroxisomal, so it wouldn’t participate in the mitochondrial NADH shuttle. However, there is experimental evidence that shows that Mdh1p forms complexes with mitochondrial enzymes of the TCA cycle and other dehydrogenases, and it has been suggested that these complexes facilitate NADH channeling. On the other hand, experiments show that in yeasts grown to saturation and stationary phases, Mdh1p is expressed without its targeting signal and stays outside the mitochondria(Chan et al., 2025). In the case of *Mucor circinelloides,* the malate dehydrogenases OAD06567.1 (MUCCIDRAFT_107142), and OAD01996.1 (MUCCIDRAFT_156403) are predicted to be mainly peroxisomal, whereas OAC97858.1 (MUCCIDRAFT_150834) is predicted as having a dual location (cytosolic >> mitochondrial). In conclusion, the location of fungal malate dehydrogenases is variable and complex, depending on the organism’s growth phase, and is difficult to predict accurately. Hence their role in each fungus NADH shuttle should be determined experimentally and in the specific growth condition being studied. In view of this, we enabled our consolidated model with the components needed to use all three types of NADH shuttles.

Notice that in the malate dehydrogenases phylogenetic tree, mitochondrial and cytosolic dehydrogenases and dual location enzymes, all mingle in the same clades. On the other hand, In the case of the aspartate aminotransferases tree, the cytosolic enzymes belong to a different clade from the mitochondrial ones (Fig 2). This clade separation according to their cell location had been previously found in other organisms’ aspartate aminotransferases (Winefield et al., 1995). This is consistent with a different evolution process for the malate dehydrogenases.

### The pentose phosphate pathway (PPP)

#### Role

Both PPP and riboneogenesis share the final steps to create ribose-5-phosphate, which is part of the nucleotides structure (a key component of ribosomes). Riboneogenesis, connects with PPP to produce S 1,7 bp. However, a full PPP process reduces NADP, while riboneogenesis does not. So, during ribose-5-phosphate production, flux through the second part of the PPP is higher than the cell’s reduced NADP requirement (Clasquin et al., 2011).

#### Curation

Many models were missing the sedoheptulose-1,7-bisphosphatase reaction (S3 Table). We updated them and added this reaction to the CFCMM.

### Interlocking cycles in central carbon metabolism Citric acid cycle

#### Role

The TCA cycle lies at the center of carbon metabolism (CoM). It connects with the 2-methylcitrate cycle (propanoate metabolism), the glyoxylate cycle (GlxC), and the GABA shunt (Huang et al., 2023) (Fig 3). Along with glycolysis and the PPP, the TCA cycle exists in all life forms, meaning it developed early (Romano & Conway, 1996). Over millions of years, it changed—some enzymes replaced others, and new steps were added or removed. Parts of it even evolved into new pathways (Huynen et al., 1999), and different organisms shaped TCA in unique ways.

**Fig 3.**
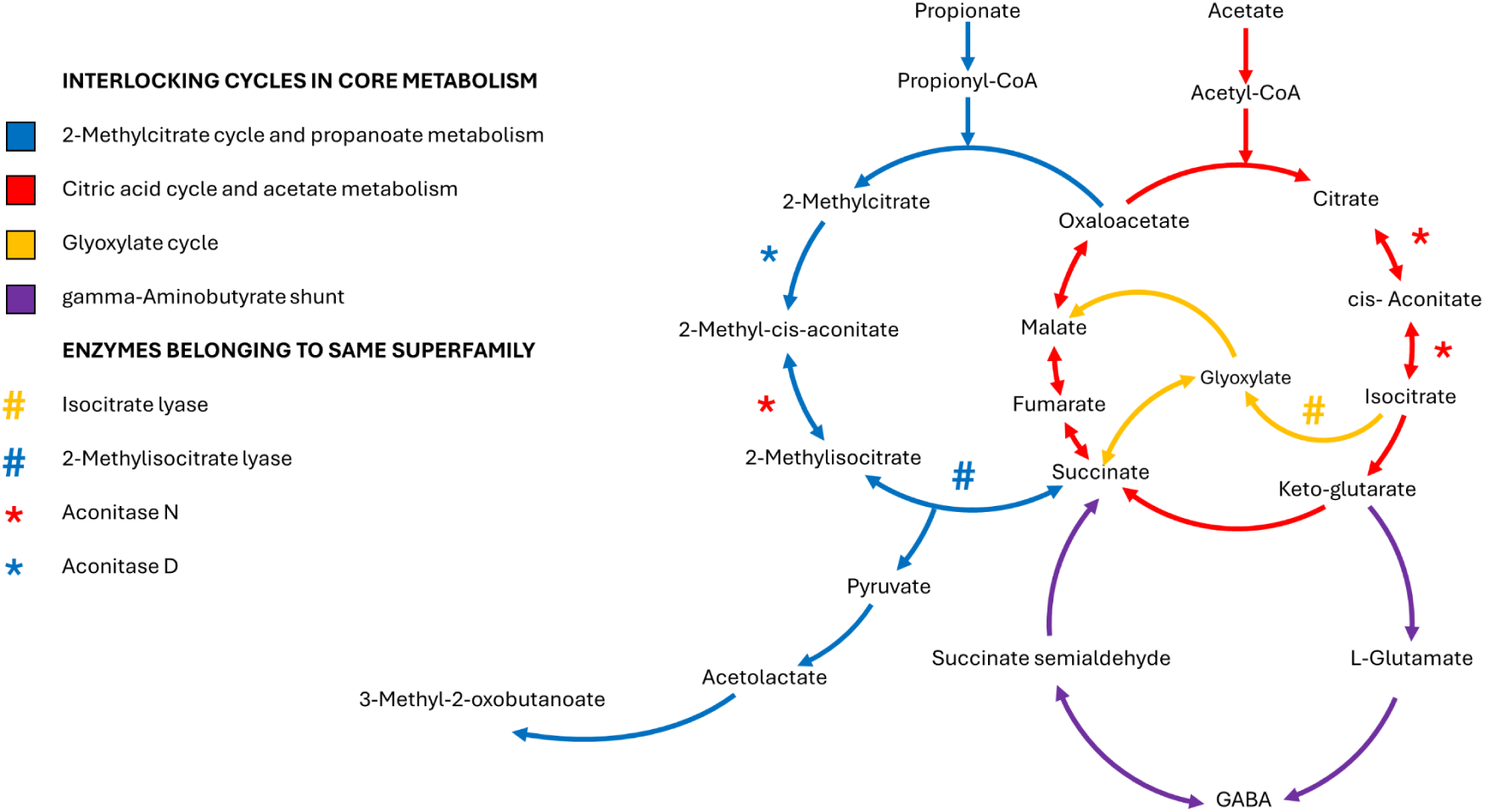
The citric acid cycle (TCA) at the epicenter of core metabolism and its interlocking with 2-methylisocitrate and glyoxylate cycles, and the GABA shunt. This is a simplification of the core metabolic network, since the TCA is mitochondrial, but the other cycles have some components located in peroxisome and cytosol. The GABA shunt components are mainly cytosolic in most fungi (S5 Fig), the glyoxylate cycle is mainly cytosolic and/or peroxisomal (S2 and S3 Figs). The 2-methylisocitrate cycle shares some enzymes with the TCA cycle (mitochondrial) and the other part of the cycle is cytosolic. Hence, the transporters for the corresponding intermediates play an important role interlocking these cycles. Notice some enzymes of the same families play similar roles in different cycles: isocitrate lyase (EC 4.1.3.1), and methylisocitrate lyase (EC 4.1.3.30); aconitate hydratase (EC 4.2.1.3), and 2-methylcitrate dehydratase (EC 4.2.1.79).

#### Curation

S3 Table summarizes variations in the TCA cycle and associated connecting reactions across the studied models. Key challenges in merging TCA cycle models were the pyruvate dehydrogenase (PyrDH) and oxoglutarate dehydrogenase (OXDH) complexes. Different methods were used to model them, leading to variations in their reactions. We organized PyrDH reactions into a set of four partial ModelSEED reactions plus one overall reaction (rxn00154_m0). Similarly, we grouped OXDH reactions into four partial ModelSEED reactions and one overall one (rxn08094_m0). Other TCA cycle reactions also needed adjustment. Some models split the two steps of aconitase (EC 4.2.1.3) into separate reactions, while others combined them. The same happened with isocitrate dehydrogenase (EC 1.1.1.42). To ensure accuracy, we included both methods in the CFCMM.

### The Glyoxylate cycle

#### Role

The glyoxylate cycle (GlxC) produces succinate (Fig 3), which moves into mitochondria and is used in the TCA cycle to make building blocks for amino acids (Kunze et al., 2006). The biggest energetic loss for the cell in running GlxC instead of the TCA cycle would be in that the conversion from isocitrate into succinate via GlxC does not produce any reduced NADH, nor ATP whereas the conversion of isocitrate into succinate via TCA cycle uses 3 reactions where not only ATP, but also 2 equivalents of NADH are produced (Each NADH gives three molecules of ATP in the ETC). The malate produced in the GlxC, can also be converted to fumarate in cytosol. There are fumarate reductases which irreversibly reduce fumarate to succinate, using NADH (Enomoto et al., 2002).

#### Curation

There is a great variation in the location of the GlxC enzymes in the models covered in this study (S7 Table and S2 and S3 Figs), but since the location of all *S. cerevisiae* enzymes has been proven in experiments (Gabay-Maskit et al., 2020; Schummer et al., 2020; Gabaldón et al., 2006; Kunze et al., 2006; Regev-Rudzki et al., 2005), we decided to use *S. cerevisiae* GlxC locations as a general guide for CFCMM. Finding and placing isocitrate lyases (rxn00336) in the models was important (S7 Table and S8 Table), to separate them from 2-methylisocitrate lyases (rxn00289), which work in propionate metabolism (S9 Table). This helped fix mistakes in the models. All isocitrate lyases belong to the same group in the AceA superfamily phylogenetic tree (S4 Fig and S8 Table) and are mainly found in the cytosol, as predicted by WoLFPSORT (Horton et al., 2007).

### The GABA shunt

#### Role

Fungi can store and use GABA as an extra source of carbon and nitrogen. Some, like *S. cerevisiae* and *A*. *nidulans*, can even survive using GABA as their only nitrogen source. The GABA shunt is a version of the TCA cycle, but instead of breaking down 2-oxoglutarate to succinate, it changes it into L-glutamate (Fig 3). Normally, 2-oxoglutarate dehydrogenase (EC 1.2.1.105) and succinate-CoA ligase (EC 6.2.1.4) would turn it into succinate through oxidation. However, taking the amination route is harder for the cell because it does not use the stored energy in succinyl-CoA (Kumar & Punekar, 1997) (Fig 3).

#### Curation

The problem of placing GABA shunt reactions in the right compartments while modeling is similar to that of the GlxC (S5 Fig). To solve this, we used the same method as for GlxC enzyme locations and based the CFCMM on the *S. cerevisiae* model. One exception is *Eremothecium gossypii*, where glutamate decarboxylase is missing from both the model and genome. In this case, GABA might be made by 4-aminobutanoate:2-oxoglutarate aminotransferase (S7 Table).

### 2-methylcitrate cycle and propanoate metabolism

#### Role

Propionyl-CoA is a harmful metabolic compound, and organisms break it down using two known pathways—one through acrylyl-CoA and the other through 2-methylcitrate (Otzen et al., 2014). Some bacteria, like *Vibrio tapetis,* use the 2-methylcitrate cycle, which is also active in pathogenic fungi (Huang et al., 2023). This cycle is part of propionate metabolism (Fig 3). *Saccharomyces cerevisiae* processes propionate into pyruvate, acetate and alanine. In *S. cerevisiae’s* Cit3p is a dual specificity citrate and methylcitrate synthase (rxn00679 in S6 Fig). In mutants lacking this activity, the addition of propionyl-CoA inhibited pyruvate dehydrogenase and caused an accumulation of acetate and isobutanol (Graybill et al., 2007). S6 Fig illustrates the complete propanoate metabolism pathway.

#### Curation

While integrating the 2-methylcitrate cycle, we found missing reactions in several models (S9 Table; Tabs A and B). To ensure accurate metabolic annotation, we checked whether these missing reactions were just gaps in certain models or if they reflected natural differences among organisms. Except for the enzyme in reaction rxn12777, all the enzymes involved in making propionyl-CoA were present in at least one model, and most had a full 2-methylcitrate pathway (S9 Table; Tabs A and B). We worked to fill in the missing steps from propionyl-CoA to 2-methyl-propanal. Using available enzyme sequences, we ran blast searches on fungal genomes where models lacked parts of the pathway. Whenever possible, we used *S. cerevisiae* sequences as reference. S9 Table; Tab C lists the proteins found through this search.

#### 2-Methylcitrate dehydratase

The data on 2-methylcitrate dehydratase (aconitase D or AcnD) in fungi models were inconsistent, since some models link AcnD to the dehydration of 2-methylcitrate and aconitase N to the hydration of 2-methyl cis-aconitate, whereas others link AcnD to both steps (S8 Table and S9 Table; Fig 4).

**Fig 4.**
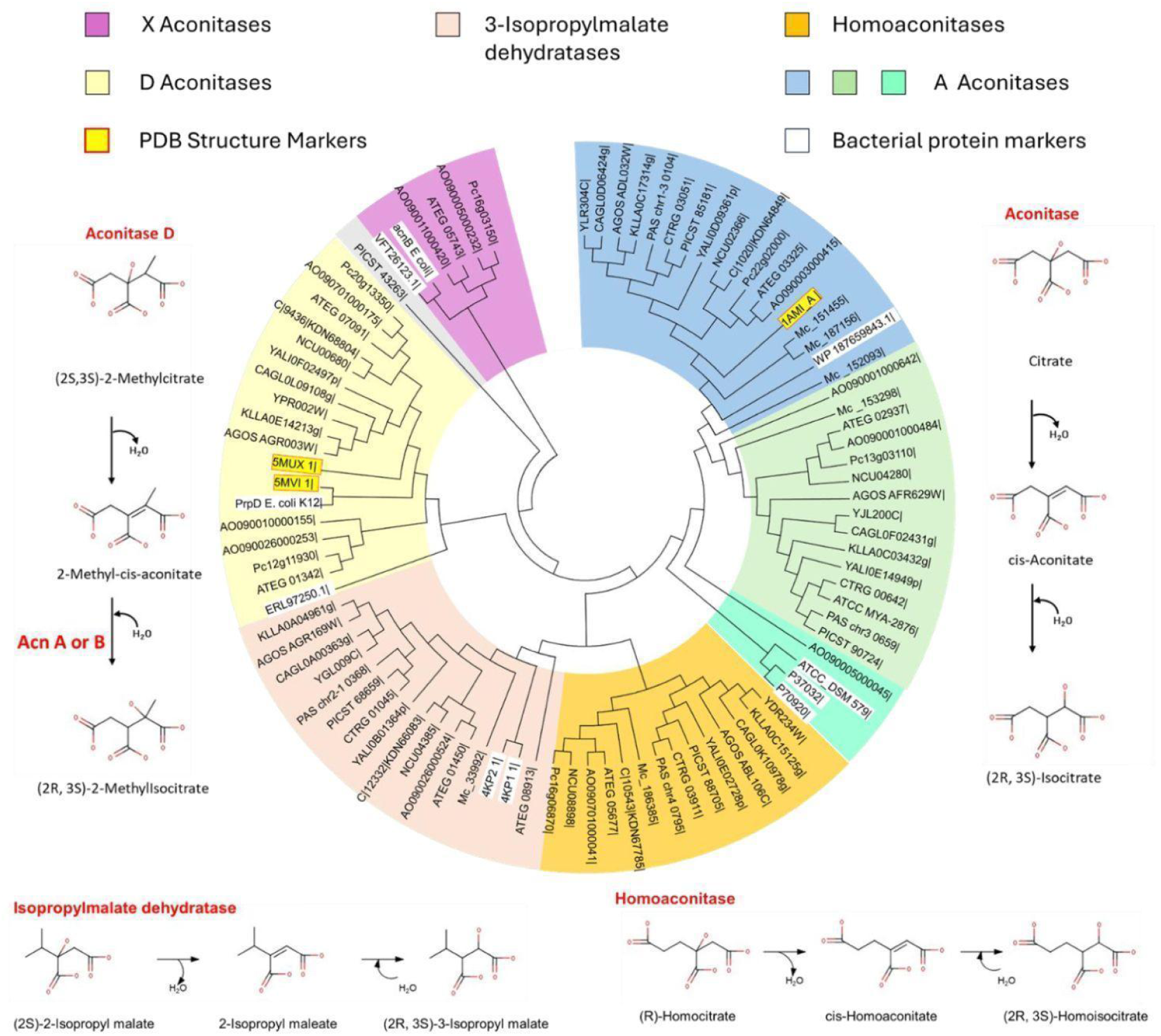
Aconitases superfamily phylogenetic tree and the reactions they catalyze. The reactions catalyzed by the different aconitase family members are shown on the margins of the figure. AcnA, AcnB, and AcnN aconitases (EC 4.2.1.3)/ 2-methylisocitrate dehydratases (EC 4.2.1.99); Homoaconitases (EC 4.2.1.36); Isopropylmalate isomerases, also called 3-isopropylmalate dehydratases (EC 4.2.1.33); AcnD aconitases called 2-methylcitrate dehydratases (EC 4.2.1.79). The fungal aconitases family tree was made using 100 replicates to infer the bootstrap consensus tree applying the maximum likelihood method (JTT matrix-based model) (highest log likelihood -15008.6359). The analysis involved 96 amino acid sequences and the evolutionary analysis was conducted in Mega6 (Tamura K et al., 2013). Reference genes are marked with bright colored tags. In turquoise blue tags the sequences corresponding to genes present in the 2-methylcitrate pathway operon (see S7 Fig). These are: AcnN and AcnD from *Burkhoderia xenovorans*LB400*, Thiomonas intermedia* K12 and *Pseudogulbenkiania ferrooxidans* 2002; In yellow tags the enzymes with crystal structures in the PDB dataset used as references, which are: 4KP1: Isoproylmalate isomerase large subunit from *Methanococcus jannaschii* (MJ0499); 4KP2: homoaconitase large subunit from *Methanococcus jannaschii* (MJ1003); 5MUX: 2-methylcitrate dehydratase (MmgE) from *Bacillus subtilis*; 5MVI: 2-methylcitrate dehydratase (PrpD) from *Salmonella enterica*; 1AMI_A: *Bos taurus* aconitase. In white tags other reference genes, which are: P37032|ACON_LEGPH: Aconitate hydratase A OS=*Legionella pneumophila subsp. pneumophila* (strain Philadelphia 1 / ATCC 33152 / DSM 7513); P70920|ACNA_BRADU: Aconitate hydratase A OS *Bradyrhizobium diazoefficiens* (strain JCM 10833 / BCRC 13528 / IAM 13628 / NBRC 14792 / USDA 110); PrpD: AAC73437.1 2-methylcitrate dehydratase, *Escherichia coli* str. K-12 substr. MG1655; VFT26123.1: aconitase B, *Pseudomonas aeruginosa*; ERL97250.1: aconitase B *Rhodobacteraceae bacterium* HIMB11; WP_252705619.1: bifunctional aconitate hydratase 2/2-methylisocitrate dehydratase *E. coli* AcnB; ATCC 27634_DSM 579: Chain A|Aconitate hydratase A|*Thermus thermophilus* (strain ATCC 27634 / DSM 579 / HB8) (300852); WP_187659843.1: aconitate hydratase, *Flavobacterium buctense*.

These mistakes happen partly because aconitase superfamily members have similar sequences but different functions, requiring more precise annotation methods. The functional issue is that AcnD cannot hydrate the aconitate intermediate to complete isomerization reaction, unlike other aconitases such as A, B, or N from the TCA cycle, homoaconitase for L-lysine biosynthesis, and isopropylmalate isomerase for L-leucine biosynthesis. Organisms solved this issue by using a TCA cycle aconitase for the second isomerization in the 2-methylisocitrate pathway (Fig 4). There is experimental evidence to support this in *Escherichia coli* (Brock et al., 2002), and in *Bacillus subtilis* (Reddick et al., 2017). There is also genetic evidence showing that these enzymes work together in bacterial 2-methylcitrate pathways. Some experiments suggest that fungi also use two different enzymes for the isomerization in the 2-methylcitrate pathway, as seen in *Yarrowia lipolytica* (Uchiyama et al., 1982). Its TCA aconitase differs in structure from bacterial AcnB, and researchers think this aconitase coordination may also occur in pathogenic fungi (Huang et al., 2023). Additionally, bacterial TCA aconitases that share structural homology with fungal aconitases, including those from *S. cerevisiae* and *E. gossypii* (Fig 4), are found in operons, where genes belonging to the TCA cycle and propionate metabolism are neighbors (S7 Fig).

#### 2-Methylisocitrate lyase

In many fungi, 2-methylisocitrate lyases (2-methylcitrate cycle/ propanoate metabolism) and isocitrate lyases (GlxC) belong to the AceA superfamily but in bacteria like *Escherichia coli,* there are methylisocitrate lyases belonging to the PrpB superfamily. However, it is thought that most fungi lost this bacterial type of methylisocitrate lyase, and after a gene duplication of the AceA isocitrate lyase, the methylisocitrate lyase subsequently evolved from it (Huang et al., 2023). Very few fungi retain some enzymes belonging to the PrpB superfamily, and these enzymes are currently linked to an oxaloacetase (EC 3.7.1.1) reaction in *A. oryzae*, *A. terreus* and *Penicillium* models (S8 Table, lyases Tab). A phylogenetic tree of the AceA and PrpB superfamilies (S4 Fig) places all the AceA 2-methylisocitrate lyases in one clade, and the WoLFPSORT program (Horton et al., 2007) locates all of them preferably in mitochondria facilitating annotation corrections (S9 Table, Tab C). Probably the fungal oxaloacetases evolved from the bacterial PrpB methylisocitrate lyases (Müller et al., 2011).

### Metabolism of sugars, fermentation, and the Crabtree effect

#### Role

Some extreme Crabtree negative yeasts can ferment glucose but either cannot grow or can grow for only a few generations using fermentation alone. As our core models demonstrate energy yields in both anaerobic and aerobic conditions, it must be flexible enough to accommodate anaerobic fermentation and aerobic respiration in these fungi. Many fungi ferment a variety of sugars, so we incorporated sugar-degradation reactions from all models included in this study. Even under anaerobic conditions, some yeasts produce metabolites in addition to ethanol, including polyalcohols and mono-, di-, and tricarboxylic acids. We therefore incorporated these reactions into the CFCMM. *Saccharomyces cerevisiae* and other Crabtree positive yeasts need certain growth factors when oxygen is low, including vitamins (nicotinate), sterols, and unsaturated fatty acids, since oxygen is required to make them. These factors must be considered when designing growth media for metabolic modeling.

#### Curation

Our CFCMM integrates models of both Crabtree positive and Crabtree negative yeasts (Table 1, S1 Table). Most of the models covered alcoholic and acetate fermentation but acetic ester fermentation reactions were present only in the *S. cerevisiae* model, and alcohol acetyltransferase only found in *S. cerevisiae* and *Candida Glabrata* genomes. On the other hand, *S. cerevisiae, Aspergillus oryzae, Kluyveromyces lactis, Scheffersomyces stipitis and Candida glabrata* are known to produce acetoin (Su et al., 2015; S. Li et al., 2015; Jiang, 1993). Probably *Aspergillus terreus, Candida Tropicalis, and Penicillium chrysogenum Wisconsin* also produce it, since they have good homologs to (R,R)-Butane-2,3-diol:NAD+ oxidoreductase. Some of the reactions related to acetoin fermentation were already present in *N. crassa, C. glabrata, C. tropicalis and S.cerevisiae* models (S10 Table). The same enzyme, which belongs to a butanediol dehydrogenase (DH) like superfamily (cd08233), reduces diacetyl to acetoin and the latter to butanediol (Heidlas & Tressl, 1990).

To complete the fermentation section in our CFCMM (Fig 5), we identified genes linked to fermentation using blast searches on known superfamilies (S10 Table). Crabtree positive and Crabtree negative yeasts can control sugar movement in different ways, and this can lead to more marked differences in ATP and reducing power between fermentation and respiration (van Dijken et al., 1986).

**Fig 5.**
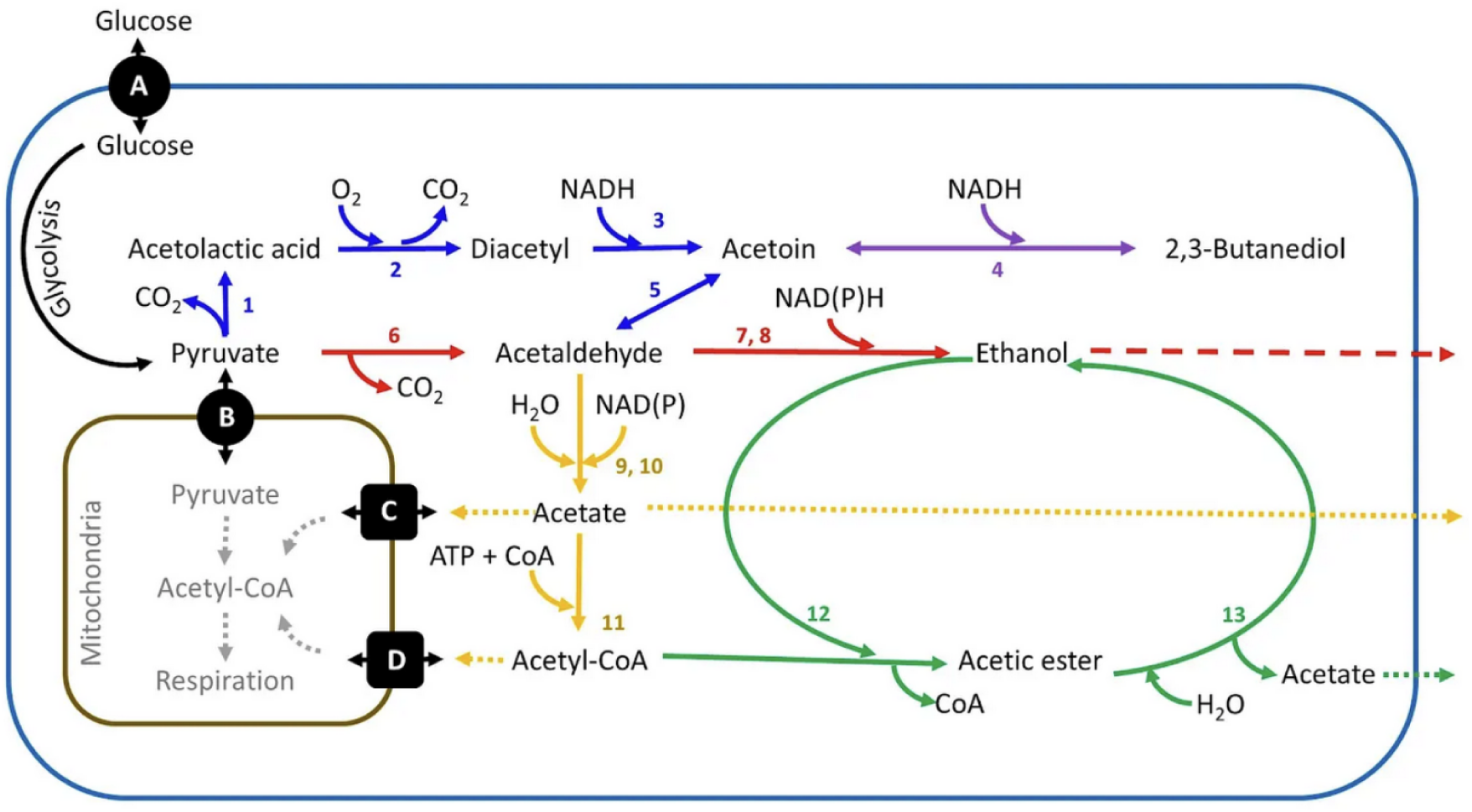
Pyruvate metabolism and fermentation. The pathways are color coded: alcoholic fermentation (red); acetoin production (blue); butanediol production (purple); acetate fermentation (golden); acetic ester fermentation (green). Notice that the diacetyl formation is a spontaneous aerobic reaction of acetolactic acid, whereas the ethanol, acetate and acetoin productions via acetaldehyde are anaerobic fermentations. Enzymes: 1, acetolactate synthase (EC 2.2.1.6); 2, spontaneous reaction; 3, acetoin: NAD+ oxidoreductase (EC 1.1.1-); 4, (R)-2,3-butanediol dehydrogenase (EC 1.1.1.4); 5, acetaldehyde condensation; 6, pyruvate carboxy-lyase (EC 4.1.1.1); 7 and 8, alcohol dehydrogenase (EC 1.1.1.2); 9 and 10, acetaldehyde:NADP+ oxidoreductase (aldehyde dehydrogenase) (EC 1.2.1.3); 11, acetate:CoA ligase (EC 6.2.1.1 or EC 6.2.1.13); 12, alcohol O-acetyltransferase (EC 2.3.1.84); 13, ethyl acetate hydrolase (EC 3.1.1.113). Transporters: A, glucose transport (diffusion); B, pyruvate proton symporter; C, acetate transport (diffusion); D, acetyl-CoA mitochondrial transport. This diagram shows their different fermentation types. One key compound, acetolactate, reacts with oxygen to make diacetyl, which enzymes can turn into acetoin. More diacetyl forms when valine levels rise. Acetolactate also breaks down without oxygen using flavin, nicotinamide or quinone as electron acceptors (KEGG database) producing acetoin (Krogerus & Gibson, 2013). The fermentation reactions, equations and definitions can be found in S10 Table.

### Construction of the Consolidated Fungi Core Metabolism Model

The 13 corrected and completed core metabolism networks were integrated into a consolidated model template (S11 Table), by incorporating all associated gene-protein-reactions (GPRs) relationships from the original models to represent each enzymatic function as comprehensively as possible. We manually added transporter reactions to ensure a functional model, particularly for transport processes in which GPR associations are traditionally difficult to identify. Certain enzymes located on the inner mitochondrial membrane and oriented toward the intermembrane space posed additional annotation challenges. These enzymes interface with both hydrophilic metabolites in the intermembrane space and hydrophobic components such as the mitochondrial ubiquinone pool. Notable examples include mitochondrial glycerol 3-phosphate dehydrogenase (Gut2) and Complex III of the respiratory chain.

Across different models, the compartmentalization of these reactions and metabolites varies—some are annotated as mitochondrial, whereas others are annotated as cytosolic—leading to inconsistencies and redundancies during model integration. To address this issue, and in alignment with the compartment definitions used in most prior models, we standardized the following assignments: (1) the ubiquinone pool was designated as mitochondrial, (2) glycerol 3-phosphate in the intermembrane space was treated as cytosolic, and (3) cytochrome c was considered mitochondrial. We then mapped the relevant enzymes from the 13 models to these standardized compartments to ensure consistency in localization and reaction representation.

During this integration process, we encountered instances of inaccurate biochemical representations, including incorrect reaction stoichiometries, redox imbalances, and inconsistencies in transport systems. These issues were systematically reviewed and corrected based on available literature and established biochemistry, ensuring that the integrated template accurately reflected biologically valid metabolic functions (S4 Table). Where necessary, we modified the corresponding existing ModelSEED reactions to match the reactions in the source models by substituting one of the substrates or products with a chemically equivalent compound (for an example, replacing ubiquinone-8 with ubiquinone-6). Each compound of these reactions was carefully mapped to corresponding ModelSEED compounds and subsequent reactions.

For reactions and their corresponding genes that were absent from all models but were required to fill pathway gaps, we introduced candidate assignments only when there was evidence that (1) the fungal enzyme catalyzes the specific reaction in question, or (2) the enzyme is known to occur in fungi even if its exact substrate specificity has not been experimentally resolved. Greater caution was applied when the enzyme was characterized only in other organisms and a fungal homolog was present without direct evidence supporting the same activity. If both the reaction and the corresponding gene were absent, or if only a very small number of fungal examples were available, we considered the signal potentially due to contamination and did not fill the gap.

### Comparative analysis of fungi core metabolism bioenergetic functions

One of the principal applications of core metabolism models is the estimation of ATP yields. Here, we demonstrate the functionality of the CFCMM by performing these calculations and comparing the results with experimental observations reported in the literature.

### Energy yield predictions for respiration

#### Role

Among fungal species, variation in ATP yield across core metabolic models is primarily driven by differences in the fungal ETC architecture, particularly the presence or absence of Complex I (Fig 6 and Table 2). In fungi that retain it, Complex I plays a dual role in their ETCs—as it donates electrons to the ubiquinone pool but also translocates protons across the inner mitochondrial membrane, thereby contributing to the proton motive force. As a result, fungal species that retain Complex I (Fig 6 and Table 2) establish a stronger proton gradient and consequently synthesize more ATP via ATP synthase, compared to fungi that have evolved ETCs lacking Complex I (Fig 6 and Table 2).

**Fig 6.**
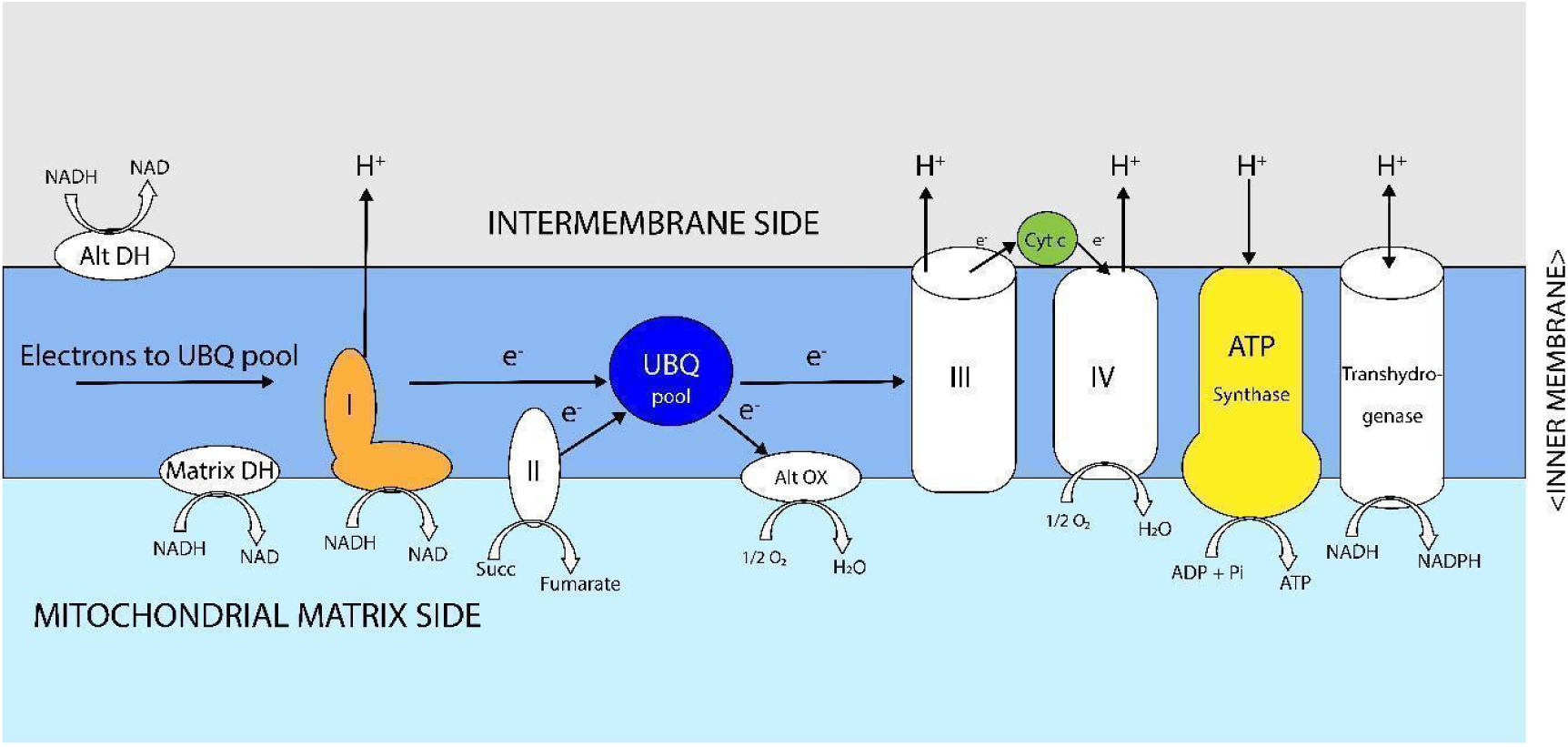
Electron transport chain (ETC) components of fungi mitochondria. Complex I, matrix and alternative dehydrogenases (Table 2 and S5 Table) donate electrons to the ubiquinone pool. Abbreviations: Matrix DH, matrix NADH dehydrogenase; Alt OX, alternative oxidase; Alt DH, Alternative NADH dehydrogenase; UBQ, ubiquinone; Cyt c, cytochrome c; Succ, succinate. In the figure the Complexes I, II, III and IV are simply represented by their Roman numbers.

#### Results

In our models’ simulations, fungi possessing Complex I achieved ATP yields of approximately 30 mmol ATP per mmol of glucose. In contrast, fungi lacking Complex I produced approximately 22 mmol ATP per mmol of glucose. This predicted difference in ATP yield highlights the central role of Complex I in enhancing energy production through oxidative phosphorylation. Under anaerobic conditions, where the ETC is inactive, all models consistently predicted an ATP yield of approximately 2 mmol ATP per mmol of glucose, generated via fermentation with ethanol as the major by-product (Table 2).

### Alternative oxidase

#### Role

Unlike the canonical ETC route, this alternative pathway reduces proton translocation across the inner mitochondrial membrane by bypassing Complexes III and IV; under these conditions, the major proton-pumping contribution comes from Complex I, when present, thereby sustaining the proton electrochemical gradient required for ATP synthesis by ATP synthase. Despite its lower energetic efficiency, AOX plays a critical role in maintaining redox homeostasis, particularly under environmental or metabolic stress. This feature is especially relevant in pathogenic fungi such as *Candida albicans*, in which AOX supports survival and metabolic activity in host environments where ETC components may be targeted by the immune system or by antifungal agents (Liu et al., 2023). By diverting electron flow away from Complexes III and IV, AOX prevents over-reduction of the ubiquinone pool, limits excessive ATP production through ATP synthase, and reduces the risk of reactive oxygen species formation, which often originates at these complexes when electron flow is disrupted (Siedow & Umbach, 2000).

#### Results

In our simulations, we evaluated ATP yields in fungal models simulating AOX function, by blocking CIII and CIV function, both in the presence and absence of Complex I. When Complex I is present, it contributes to proton translocation across the mitochondrial membrane, and electrons subsequently reduce oxygen via AOX. Under these conditions, the model predicts an ATP yield of approximately 12 mmol ATP per mmol glucose. In contrast, in models of organisms lacking Complex I, blockage of CIII and CIV causes a substantial ATP yield drop to 4 mmol ATP per mmol glucose (Table 2). These functional analysis using our models aligns with previous reports that AOX respiration is energetically inefficient due to its poor proton translocation in the ETC, and that the presence of Complex I or other proton-pumping components significantly impacts ATP output(Joseph-Horne et al., 2001; Siedow & Umbach, 2000; Vanlerberghe & McIntosh, 1997).

### ATP budget for fermentation

#### Role

Some fungi cannot survive using fermentation alone as their sole source of energy. Although they ferment glucose under low-oxygen conditions (De Deken, 1966), they remain oxygen dependent, and petite mutants are lethal in these organisms (van Dijken et al., 1986). Fermentation yields relatively little energy, at most 2 mol ATP per mol glucose through glycolysis. The overall glycolytic equation is as follows: Glucose + 2 ADP + 2Pi + 2 NAD => 2 pyruvate + 2 ATP + 2 H2O + 2 NADH + 2 H+

#### Results

The metabolic simulations in all our models were consistent with these calculations, predicting an ATP yield of approximately 2 mmol ATP per mmol of glucose, generated via fermentation under anaerobic conditions and producing ethanol as a by-product (Table 2).

Beyond respiration, our central carbon core models also highlight the importance of NADH shuttle systems in maintaining redox balance and linking cytosolic and mitochondrial metabolism. These shuttles provide fungi with metabolic flexibility, especially in species lacking Complex I, and help explain differences in energy efficiency across diverse taxa.

## Conclusions

Reconciling fungal core metabolism into a unified biochemical framework is a prerequisite for rigorous cross-species comparisons and provides a scalable foundation for accurate GEM reconstruction. By standardizing reactions, cofactors, and compartment assignments across diverse fungi, we establish a shared biochemical space in which metabolic capabilities can be examined on common footing and evaluated systematically. This harmonization resolves common inconsistencies in legacy models and enables the construction of core models for any sequenced fungal genome with greater confidence in reaction identity, mass/charge balance, and localization fidelity.

Prioritizing core metabolism, the conserved network underpinning energy generation, redox homeostasis, and precursor biosynthesis—provides a stable foundation for genome-scale metabolic modeling. Because core pathways govern stoichiometry, cofactor coupling, and baseline flux distributions, inconsistencies at this level propagate downstream and can distort pathway interpretation, gap-filling, and predicted phenotypes. A reconciled core therefore reduces uncertainty, improves reproducibility and offers a principled starting point for subsequent genome-scale reconstruction.

In summary, our reconciliation effort delivers both an enabling resource and a practical workflow for fungal systems biology: a standardized core template that supports automated draft reconstruction, principled extension to full GEMs, and a biochemically consistent substrate for subsequent comparative analyses of fungal bioenergetics. By making this framework deployable in KBase, we aim to accelerate community-wide model generation and refinement, and to strengthen mechanistic insight into how fungal metabolism varies across phylogeny, lifestyle, and environment.

## Examples of practical applications

Since core metabolism provides the cell with energy, redox equivalents and building blocks (Noor et al., 2010), changes in it can be either markedly advantageous or deleterious to the cell and its analysis has greater impact on fungal biology studies. Moreover, many fungal CoM compounds are industrial commodities themselves, or they are their close precursors. Hence, one application relevant to metabolic engineering, is the use of CFCMM in simulations to explore these metabolites production optimization. For example, some *Aspergillus* species are industrial producers of TCA cycle metabolites, and they can also be used, as well as acetyl-CoA and amino acids to synthesize many secondary metabolites (Chroumpi et al., 2020). Changes in edible fungi CoM, like in *Hypsizygus marmoreus*, can increase their production (Lin et al., 2023). Hence, edible fungi CFCMM could be useful in the design of transformants with higher sugar assimilation rates that would result in faster growth. On the other hand, simulations using pathogens’ CFCMM in combination with comparative genomics could be helpful in the production of medicines and fungicides that target CoM functions. Such is the case o*f Candida albicans* glyoxylate cycle’s isocitrate lyase and malate synthase1; gluconeogenic pathway’s PEP carboxykinase and sugar metabolism’s trehalose 6-phosphate synthases 1 and 2, which do not have homologs in humans but whose function is crucial for this pathogen’s virulence (Wijnants et al., 2022).

## Future perspectives

Although the construction of GEMs is more complex than that of CoM models, our next step is to expand the highly curated CFCMM by incorporating lateral pathways and thereby complete a comprehensive fungal template for GEM development. To support this objective, we have translated the existing fungal GEMs used in this study into the ModelSEED ontology, thereby establishing a standardized framework for model integration and extension.

With the CFCMM now completed, a substantial portion of this effort is already in place. Given the rapidly growing number of sequenced fungal genomes (https://1000.fungalgenomes.org/), the ability to streamline the generation of fungal GEMs, particularly those with accurately curated energy metabolism, will substantially accelerate research across the broader fungal science community.

## ABBREVIATIONS IN THE TEXT

DH: Dehydrogenase
DOE: Department of Energy
EA: Ethanol acetaldehyde shuttle
EC: Enzyme Commission
ETC: Electron transport chain
FBA: Flux Balance Analysis
G3P: Glycerol 3-phosphate
G3PS: Glycerol 3-phosphate shuttle
GABA: gamma-Aminobutyrate
GEM: Genome scale metabolic model
GlxC: Glyoxylate cycle
GPR: Gene protein reaction (associations)
KBase: Knowledge database
KO: KEGG Orthology
MAS: Malate-aspartate shuttle
MoS: ModelSEED
OXDH: Oxoglutarate dehydrogenase
PDB: Protein Database
PEP: Phosphoenolpyruvate
PPP: Pentose phosphate pathway
PyrDH: Pyruvate dehydrogenase
TCA: Tricarboxylic acid cycle.

(The rest of the abbreviations are common knowledge. Abbreviations used in the figures are given in the corresponding figure legends)

## Acknowledgements

This work is supported as part of the BER BRaVE project KP1601011 and by KBase. KBase and BRaVE are funded by the U.S. Department of Energy, Office of Science, Office of Biological and Environmental Research under Award Numbers DE-AC02-06CH11357, and DE-AC02-98CH10886.

The submitted manuscript has been created by UChicago Argonne, LLC, Operator of Argonne National Laboratory (“Argonne”). Argonne, a U.S. Department of Energy Office of Science laboratory, is operated under Contract No. DE-AC02-06CH11357. The U.S. Government retains for itself, and others acting on its behalf, a paid-up nonexclusive, irrevocable worldwide license in said article to reproduce, prepare derivative works, distribute copies to the public, and perform publicly and display publicly, by or on behalf of the Government. The Department of Energy will provide public access to these results of federally sponsored research in accordance with the DOE Public Access Plan. http://energy.gov/downloads/doe-public-access-plan

## Integral Modeling and Analysis of Fungal Core Metabolism

Janaka N Edirisinghe et al., 2026

Supplementary material; figures

**S1 Fig.**
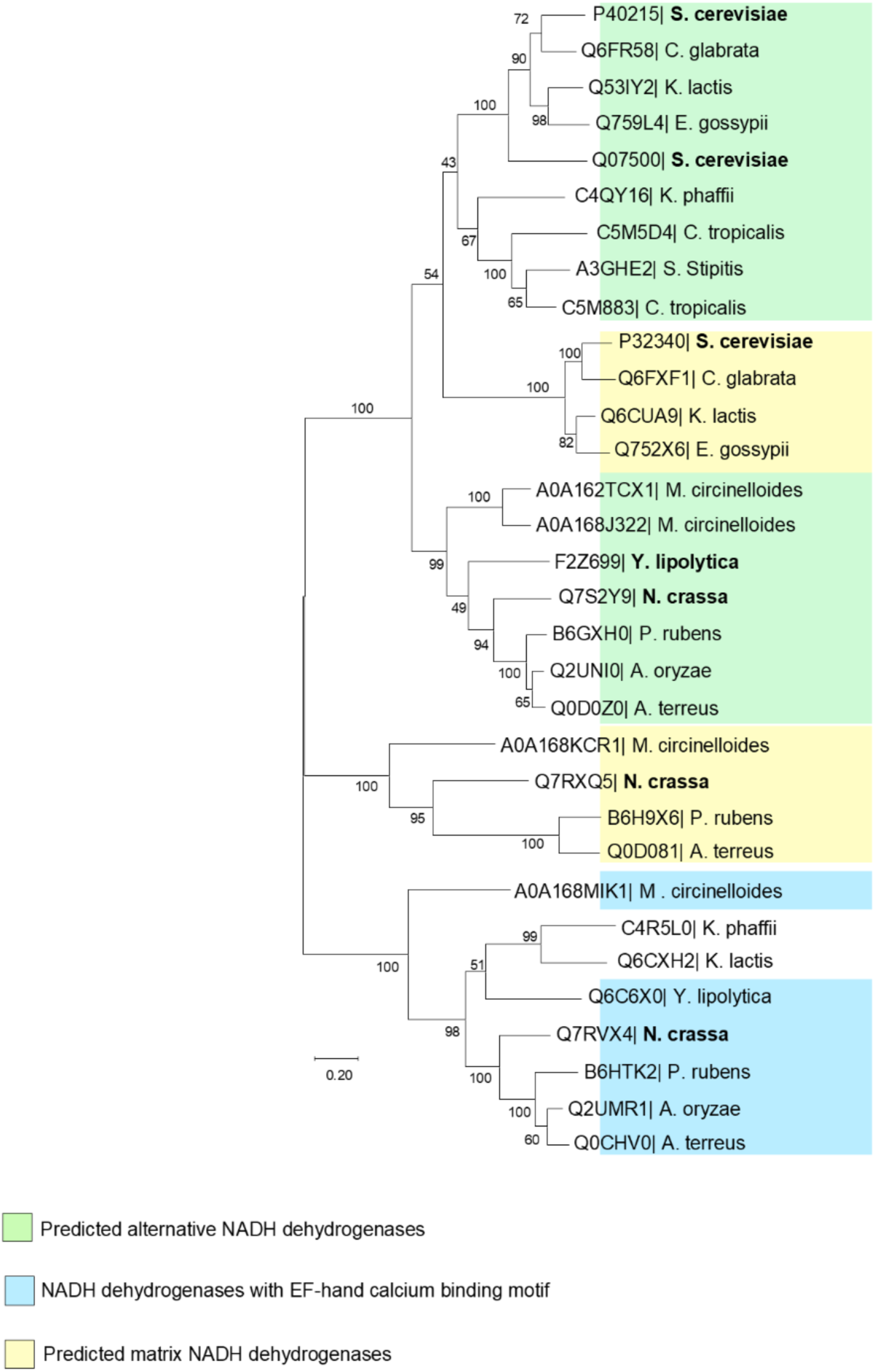
Phylogenetic tree of fungal mitochondrial NADH dehydrogenases. Enzymes identified by their UniProt numbers. Tree obtained using the MEGA6 program, clustal alignment, maximum likelihood tree. This tree shows duplication of mitochondrial alternative NADH dehydrogenases, which then would allow location of the enzymes on both sides of the organelle’s inner membrane providing an advantage for yeasts growing in anaerobic environments and using fermentative metabolism. Bold letters indicate that the enzyme locations have been determined experimentally (see text for details). A rough prediction of their position on the mitochondrial membrane has been attempted based on the experimentally proved position of their neighbors in the clade. Most models of fungi that lack Complex I allocate NADH dehydrogenase in both alternative and matrix positions, whereas models of fungi with Complex I vary in their allocation (see Table 2).

**S2 Fig.**
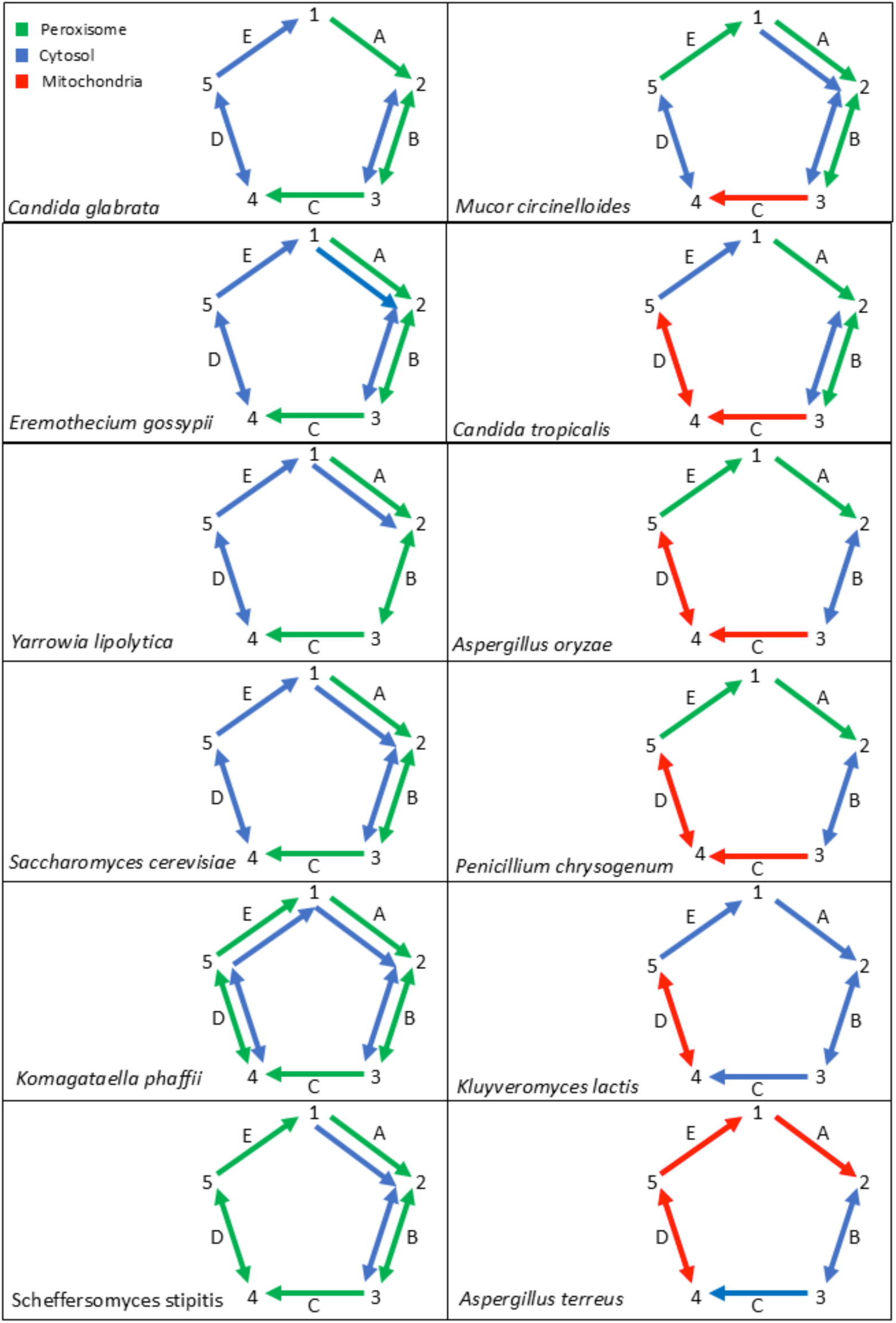
Variety of compartmentation of glyoxylate pathway reactions in the fungi models considered for the consolidation plan. Reactions’ locations color codes as shown on the top left panel. The enzymes shown here are as follows: A, malate synthase; B, malate dehydrogenase; C, citrate synthase; D, aconitase; E, isocitrate lyase. Metabolites are: 1, glyoxylate; 2, (S) malate; 3, oxaloacetate; 4, citrate; 5, D-threo-isocitrate (often abbreviated as isocitrate). Notice the absence of peroxisomal reactions in A. terreus and K. lactis. These models lack a peroxisome compartment.

**S3 Fig.**
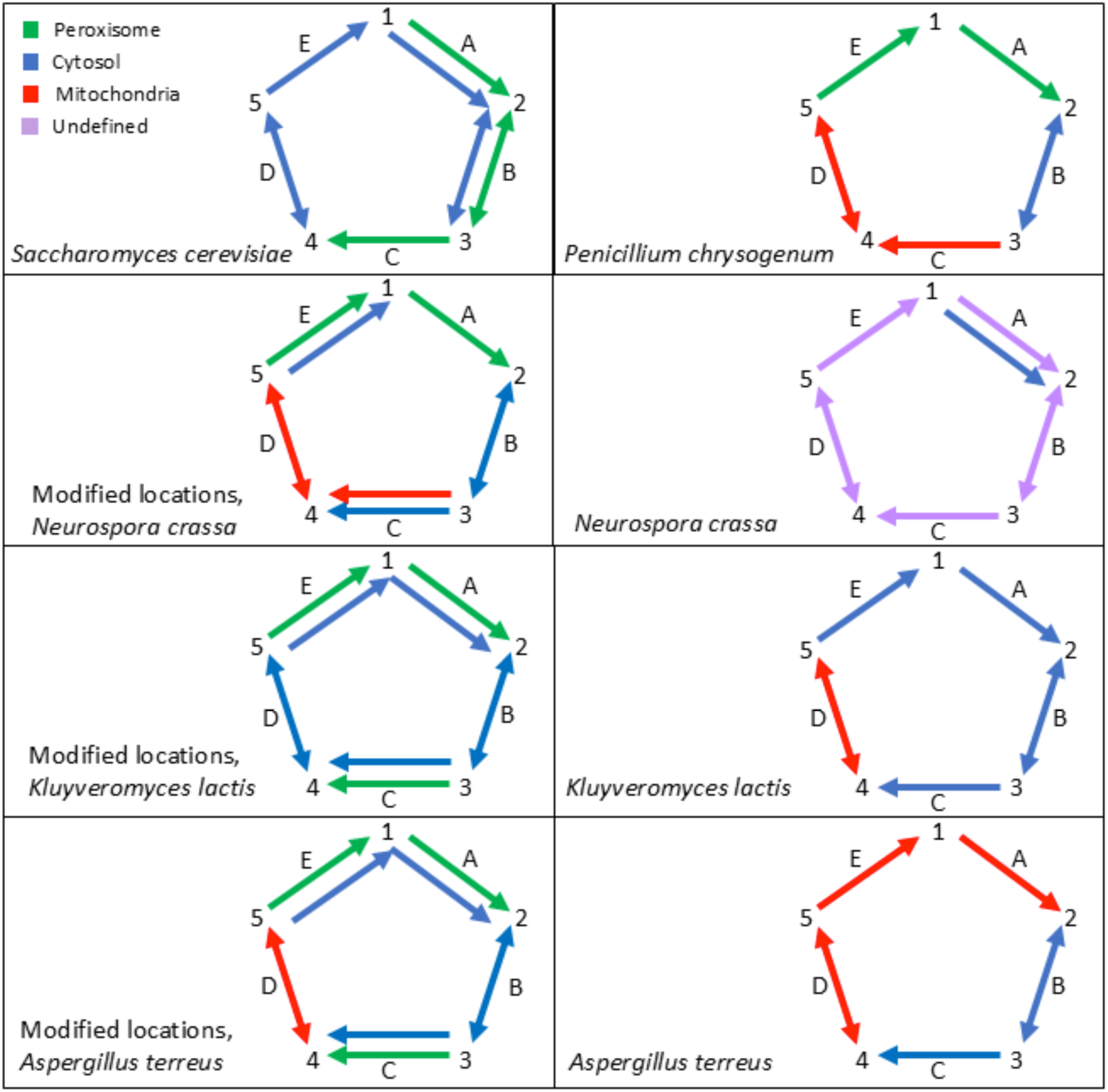
Suggestion for glyoxylate cycle reactions’ location for models which originally lack the peroxisome compartment. Top left panel: Reactions’ locations color codes shown as a reference. Enzymes and metabolites representations are the same as in the previous figure. Top panels show *S. cerevisiae* and *P. chrysogenum* location patterns as references. The locations of the enzymes shown in the modified glyoxylate pathway for *N. crassa, K. lactis*, and *A. terreus* are predicted using the WoLFPSORT program as described in S7 Table.

**S4 Fig.**
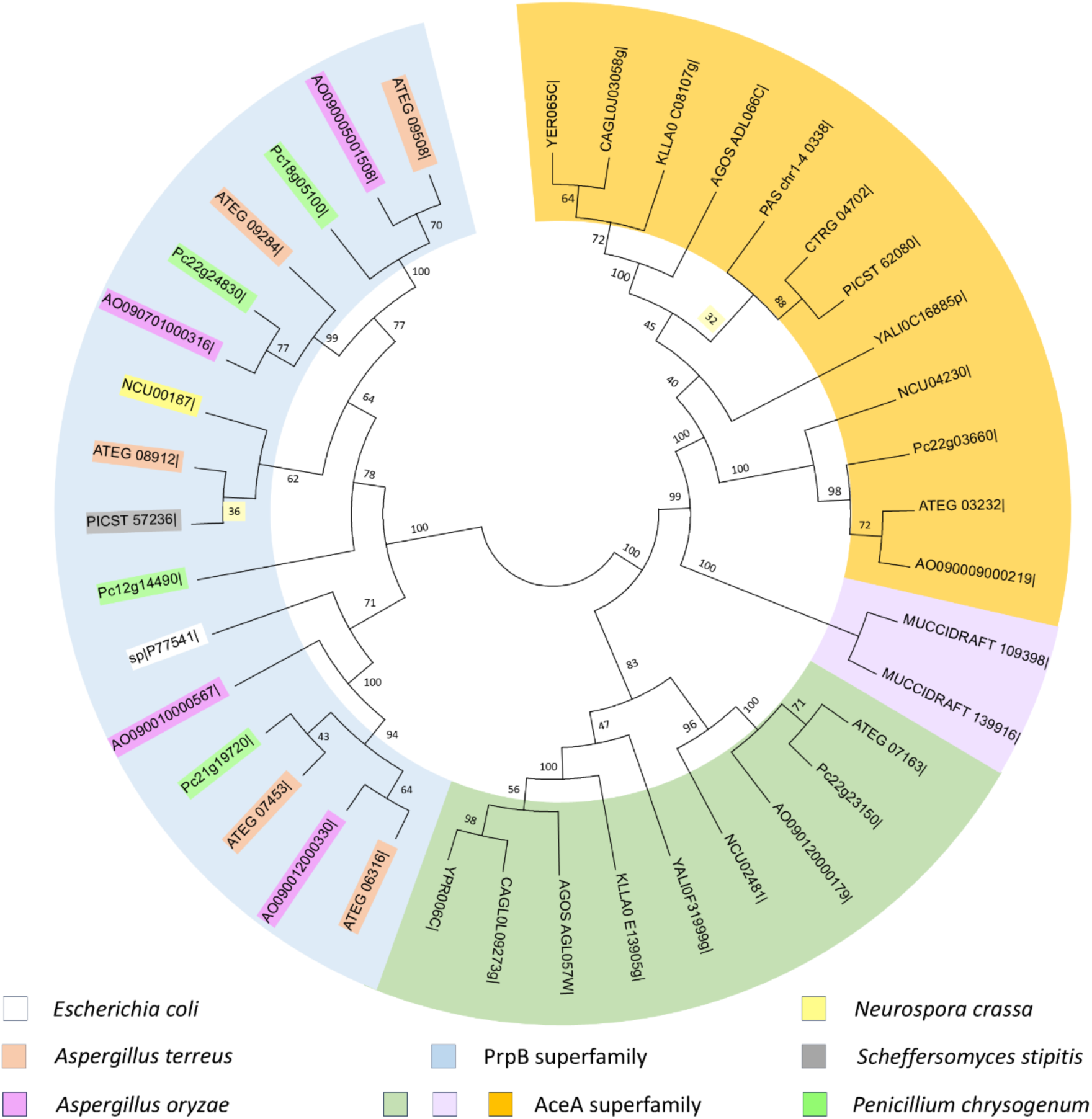
Phylogenetic tree of fungal PrpB and AceA lyases. Most fungi in our study have methylisocitrate or isocitrate lyases belonging to the AceA superfamily, but only 5 of them also have lyases belonging to the PrpB superfamily. AceA methylisocitrate lyases clade highlighted olive green. AceA isocitrate lyases clades highlighted in orange and lavender. The *Mucor circinelloides* has two isocitrate lyases of the AceA superfamily in a separate clade. Each enzyme in the PrpB superfamily clade is color-coded according to its source. The *Escherichia coli* PrpB methylisocitrate lyase is used as a marker. Protein sequences obtained from https://narrative.kbase.us/narrative/49164. Evolutionary history analyzed using the MEGA6 program https://www.megasoftware.net/. Clustal alignment. Bootstrap consensus tree inferred from 100 replicates. Maximum Likelihood tree with highest log likelihood of -10965.8507.

**S5 Fig.**
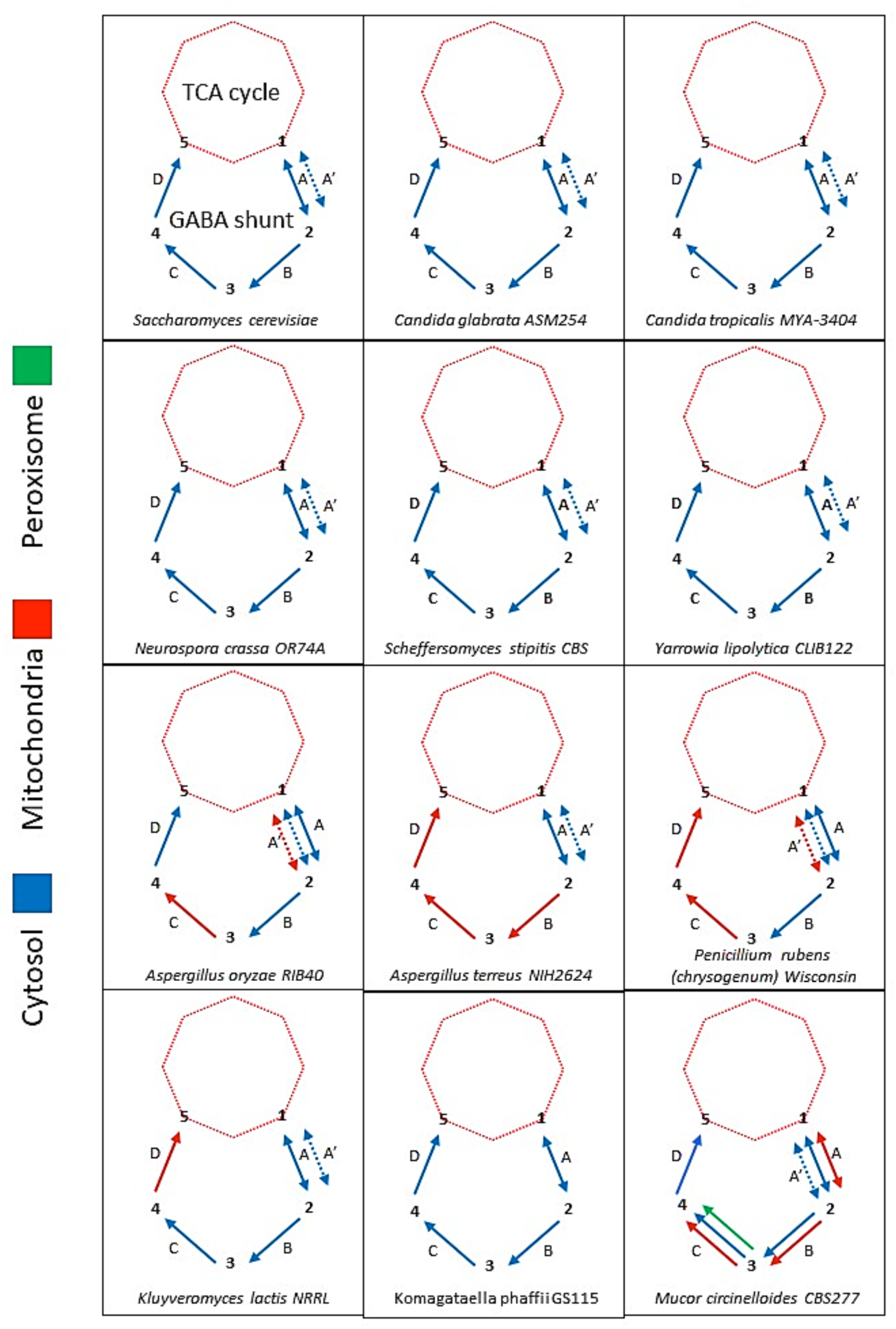
Compartmentation of the GABA shunt reactions in the fungi models used for the consolidation plan. Location color codes as shown on the left margin. The enzymes are as follow: A, glutamate dehydrogenase (NADP+) (EC 1.4.1.4); A’, glutamate dehydrogenase (NAD) (EC 1.4.1.2); B, glutamate decarboxylase (EC 4.1.1.15); C, 4-aminobutyrate-2-oxoglutarate transaminase (EC 2.6.1.19); D, succinate-semialdehyde dehydrogenase (NAD(P)+) (EC 1.2.1.16) and succinate-semialdehyde dehydrogenase (NADP+) (EC 1.2.1.79). Metabolites are: 1, 2-oxoglutarate; 2, L-glutamate; 3, gamma-aminobutyrate (GABA); 4, 4-oxobutanoate, succinate semialdehyde; 5, succinate. *Eremothecium gossypii* model’s GABA shunt is incomplete, for location of reactions see S7 Table. The metabolites 2-oxoglutarate and succinate are shared by the TCA cycle and the GABA shunt. All TCA enzymes are in mitochondria and metabolites need to be transported to the locations where the GABA shunt enzymes are located.

**S6 Fig.**
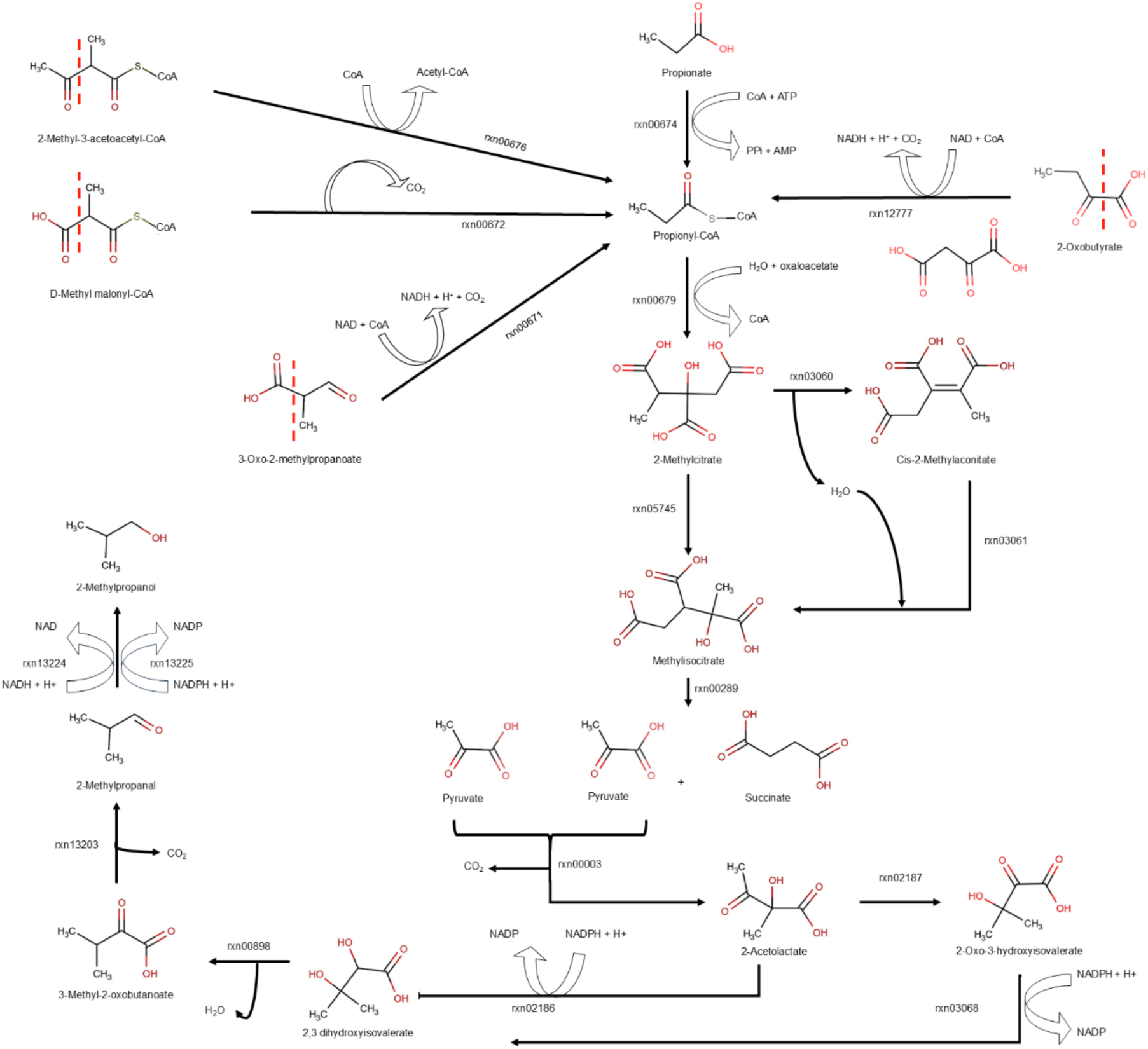
Integration of the propanoate metabolism of the fungi models. Figure shows structures of the propionyl-CoA precursors with a dash line separating the propionyl and propionyl-CoA moieties in them. Several enzymes produce propionyl-CoA and at least three of them are present in fungi models (S9 Table,Tabs A, B and C). One of the ways to dispose of this toxic intermediate is to convert it to 2-methylcitrate and enter the latter into the 2-methylcitrate cycle. This cycle is connected to the citric acid cycle by the 2-methylisocitrate lyase reaction which produces succinate. The other intermediate, pyruvate, can then be processed to an aldehyde, or even an alcohol. The overall aconitase reaction is divided into two partial reactions: the dehydration step and the addition of water to the double bond of methylaconitate. The reductive isomerization of 2-acetolactate into 2,3 dihydroxyisovalerate is represented by a single reaction in some models, or by an isomerization reaction, followed by a reduction in other models (for more details see S9 Table, Tab A).

**S7 Fig.**
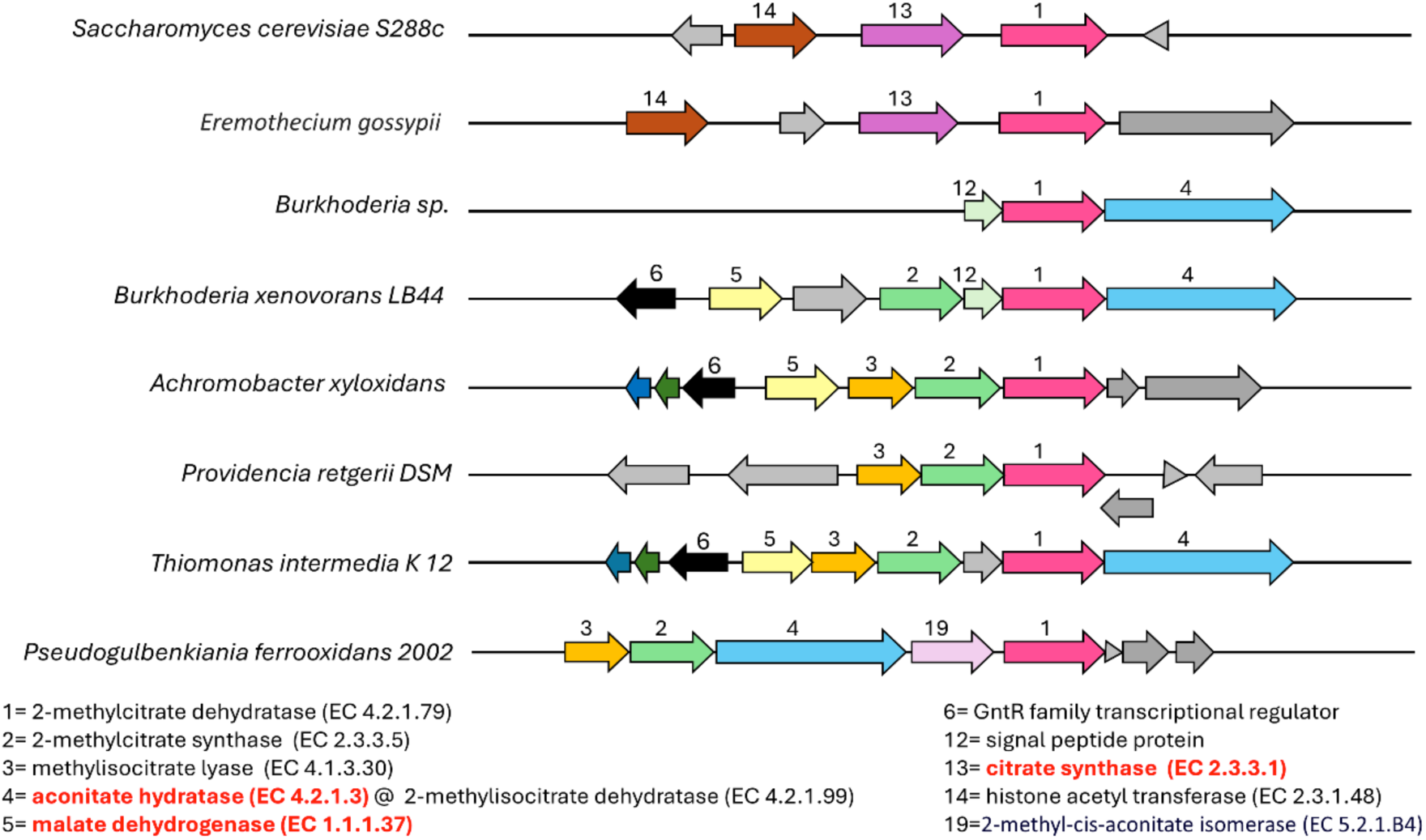
Chromosomal region of fungi and bacteria genomes containing 2-methylcitrate pathway gene(s). The regions of two fungi (*Saccharomyces cerevisiae* and *Eremothecium gossypii)* and several bacteria with the 2-methylcitrate dehydratase (EC 4.2.1.79) closest gene homologs (pink arrow [1]) at the center and surrounded by their neighboring genes. Chromosomal region viewed using the SEED database tools (https://theseed.org/wiki/Home_of_the_SEED). In the bacterial 2-methylcitrate pathway operons, this gene is accompanied not only by others encoding for this pathway’s enzymes (black type [2],[3],[5],[19]), but also by citric acid cycle genes (red bold type, [4], [5]). Interestingly, despite the absence of operons in eukaryotes, In the fungal genomes, the citrate synthase (EC 2.3.3.1) gene is next to the 2-methylcitrate dehydratase (EC 4.2.1.79) one. The ID numbers reflect the frequency with which the neighbor genes accompany the central gene, numbered [1]. The genes that are not relevant are colored grey. The 2-methylcitrate dehydratase genes (AcnD) for Burkhoderia xenovorans*, Thiomonas intermedia*, and *Pseudogulbenkiania ferrooxidans* and their aconitase/2-methylisocitrate dehydratase (AcnN) genes were used as markers in the aconitase superfamily phylogenetic tree (Figure 4).

